# SpatioEv: Spatial evolution of protein and morphological features reveals development dynamics of cells and spatial neighbourhoods

**DOI:** 10.1101/2025.06.30.662328

**Authors:** Shihong Wu, Sakina Amin, Carl Lee, Jean-Baptiste Richard, Nabeel Merali, Caroline Morrell, Lauren Overend, Weijia Gao, Felicia Tucci, Alex Gordon-Weeks, Adam Frampton, Nicola Annels, Emma Culver, Michael Dustin, Kim S. Midwood, Rachael Bashford-Rogers

## Abstract

Understanding cellular function in tissues demands sophisticated tools to decode complex microenvironmental interactions. Current spatial analysis methods often lack the comprehensive framework needed to systematically analyse cell morphology, dynamics, interactions, and extracellular matrix (ECM) architecture. We introduce SpatioEv, a unified computational framework for highly multiplexed tissue imaging that addresses these critical gaps. SpatioEv integrates automated quality control, cell phenotyping, neighbourhood identification, multi-scale spatial characterization, niche boundary analysis, ECM fiber-cell interaction mapping, and spatiotemporal trajectory inference. This pipeline enables reproducible cell annotation, reveals novel ECM-cell interactions, characterizes tissue neighbourhood boundaries, and infers developmental progressions directly from spatial data. Using this, we can identify disease-specific spatial signatures distinguishing rheumatoid arthritis from osteoarthritis, characterize diverse tumour boundary phenotypes in cancer metastases in liver, and map evolutionary trajectories in pancreatic ductal adenocarcinoma (PDAC) at single-niche resolution. Our findings highlight the significance of spatial context in shaping cell behaviour and underscore its potential to uncover emergent tissue architecture and cellular dynamics. By addressing major analytical challenges, SpatioEv provides a scalable, adaptable platform for advancing spatial biology and translational research.

## Introduction

In living tissues, cellular function is profoundly influenced by spatial context, including the location of cells, their interactions with neighbouring cells^1–3^ and extracellular matrix (ECM)^4–10^, their spatial organization^11–15^, and the mechanical forces they experience^16–19^. However, conventional analytical methods lack the necessary framework to systematically decode these complex microenvironmental interactions, where subtle differences in cell positioning and ECM architecture can have significant functional consequences. Therefore, there is a critical need for new methodologies capable of uncovering this overlooked layer of biological organization.

Advances in single-cell profiling technologies, such as single-cell RNA sequencing (scRNA-seq), single-cell assay for transposase-accessible Chromatin using sequencing (scATAC-seq), and cellular indexing of transcriptomes and epitopes by sequencing (CITE-seq), have provided unprecedented molecular resolution for characterizing cellular heterogeneity and function in diverse physiological and pathological states^20–22^. However, these powerful approaches necessitate tissue dissociation, thereby abrogating the native spatial context and precluding the analysis of critical microenvironmental relationships and tissue architecture^23,24^. Furthermore, tissue dissociation can lead to the selective loss of fragile cell types^25^ and the disruption of delicate cellular orientations and interactions vital for understanding processes such as immune cell infiltration, tumour progression, and ECM remodelling. The limited number of cells typically profiled in single-cell suspensions can hinder the comprehensive capture of rare cell populations and the full spectrum of cellular heterogeneity present within a tissue^26,27^. Notably, the ECM is a crucial structural and signalling component in numerous biological contexts^4,28^ and is typically lost or removed during tissue dissociation^29^.

Highly multiplexed imaging methods, particularly those applicable to whole-slide images (WSIs), offer a complementary strategy by enabling detailed, *in situ* characterization of cell types and states at single-cell or even subcellular resolution while preserving the native tissue architecture^13,18,30–33^. These techniques, capable of simultaneously detecting tens to hundreds of protein or nucleic acid targets across millions of cells, hold immense potential for elucidating the spatial organization of microenvironments within disease context^12,34^. However, current computational pipelines for analysing such high-dimensional spatial data, including tools like SCIMAP^35^ and TissueSchematics^36^ , often primarily focus on pairwise cell-cell interactions and neighbourhood identification^37^ . These approaches frequently encounter challenges related to accurate cell annotation^38^. Strikingly, researchers largely rely on semi-manual annotation of cell types which, whilst considered as a gold-standard, is labour-intensive, time-consuming and highly subjective, especially across large imaging datasets or tissue microarrays. The ability to automate cellular phenotyping with high confidence is critical to allow reliable downstream interpretations.

Many key aspects of tissue organisation are underexplored in current analyses. For example, the boundaries between cell clusters or neighbourhoods represent critical zones of potential cellular signalling perturbations and interactions^39–41^. Furthermore, current methodologies often fail to capture cells undergoing dynamic transitions between states^42–45^. While the physical characteristics of the ECM, such as fiber density, alignment, length, and width, are known to significantly influence cellular behaviour^4,10,46–48^, there is a notable lack of robust tools for conducting per-fiber or per-cell-level fiber-cell interaction analyses within highly multiplexed imaging datasets. Additionally, given that progressive changes in tissue architecture, such as the coexistence of precancerous and invasive lesions, can be captured on a single image or sample^49,50^, there is a lack of established methodologies for performing spatial pseudo-temporal analysis at both the cellular and neighbourhood levels to track these evolutionary changes^51,52^. Finally, while researchers routinely explore correlations between identified features, these analyses often focus on a limited set of pre-selected targets, potentially overlooking other biologically relevant, condition-correlated spatial features.

To overcome each of these limitations of current spatial analysis approaches and to enable a more comprehensive, interpretable and unbiased interrogation of highly multiplexed spatial imaging data across diverse disease contexts, we developed **SpatioEv**, a novel, unsupervised, and interpretable end-to-end Python-based framework. To enhance accessibility for researchers with limited computational expertise, we provide step-by-step Jupyter notebooks that guide users through data navigation and analysis workflows. SpatioEv encompasses (a) a rigorous automated quality control for cyclic staining and imaging, (b) unbiased automated phenotyping based on gaussian model fitting, (c) nuanced cell state inference using a probabilistic approach, (d) automated neighbourhood identification, (e) niche boundary analysis, (f) integrated spatial analysis of fiber-cell interactions, (g) spatiotemporal trajectory inference of recurrent spatial niches, and (h) an unsupervised, multi-scale feature extraction approach that supports interpretable, statistical group-wise image comparisons (**Figure 1A**). This unified pipeline addresses key analytical challenges in the field and provides a powerful platform for revealing the complex spatial architecture and dynamic cellular interactions that underpin tissue organisation, disease progression and enables a more holistic interrogation of the complex spatial relationships within tissue microenvironments.

**Figure 1.**
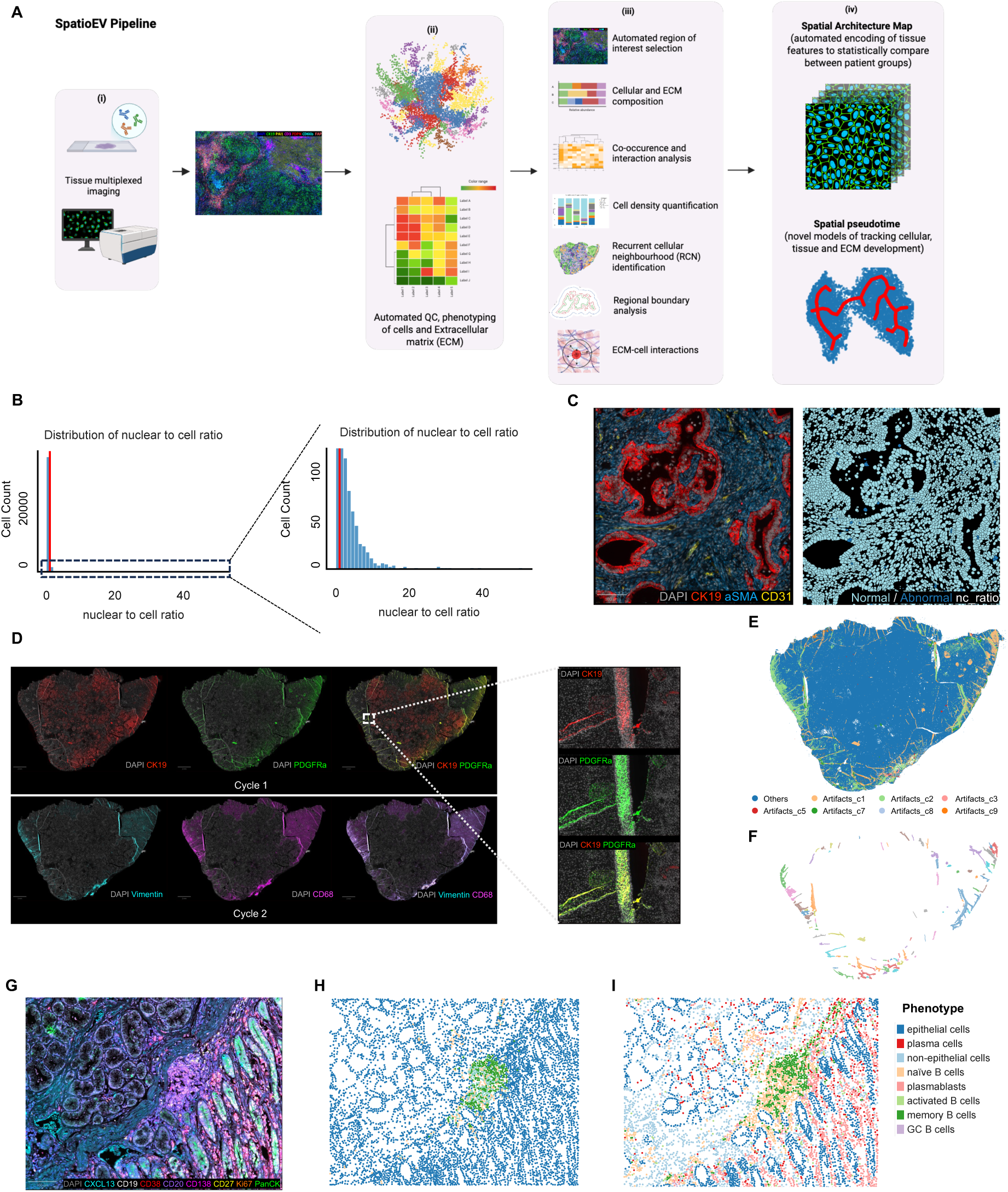
Quality control. **A.** Schematic of SpatioEv workflow. **B.** Histograms showing the distribution of Nuclear-to-cell ratio of the cell segmented on the pancreatic ductal adenocarcinoma (PDAC) image. The red line represents x=1. **C.** Representative PDAC image with a color overlay (left), followed by cell segmentation masks colored according to the nuclear-to-cell ratio (right). **D.** Representative images of cancer metastasis in liver with tissue folding artifacts (zoomed view) from cycle 1 (upper panel) and cycle 2 (bottom panel). **E.** Identification of artifact-prone cells generated from each cycle, with colors representing cycles. **F.** Artifact region removal using DBSCAN clustering based on local density patterns. **G.** Representative field of view (fov) from a PDAC image using the 9-plex B cell Panel. This panel shows a typical region of interest within the PDAC dataset. **H**. Cell annotations generated by standard SCIMAP. **I**. GMM-based cell annotations using SpatioEv.

## Results

### Novel automated robust quality control: segmentation, artifacts and phenotyping errors

First, we considered the problem of imaging quality control (QC), which includes the steps of segmentation, removal of artifacts and phenotyping (**Figure 1A-left**). Accurate cell segmentation and subsequent phenotyping remains a significant challenge in imaging data analysis due to heterogeneity of cell shape, size and density^53^, as well as the inherent limitations of representing 3-dimentional structures with 2-dimensional images^54^. To address segmentation inaccuracies, SpatioEv quantifies cellular morphological parameters, including cell area and nuclear-to-cell (N:C) area ratio, to identify and exclude biologically implausible cells (outliers). Since nuclei are typically confined within cells, aberrant N:C ratios often reflect segmentation errors or imaging artifacts. These may arise from technical factors such as overmerged nuclei due to thresholding errors (e.g., DAPI/Hematoxylin over segmentation), incorrect algorithm parameters, weak membrane staining, preprocessing artifacts (e.g. excessive smoothing), or misalignment. Biological contributors include apoptotic or necrotic cells with condensed nuclei or multinucleated cells (e.g. tumor giant cells) erroneously merged during segmentation. By quantifying the proportion of such artifacts, SpatioEv assesses image quality and excludes problematic regions to ensure robust downstream analysis. For example, in pancreatic ductal adenocarcinoma (PDAC) tissue sections, cells with N:C ratios > 1 (e.g., **Figure 1B-C, Supplementary Figure 1A**) were excluded to ensure high-quality downstream quantification of intact cells.

Multiplexed imaging of whole-slide tissue sections frequently introduces artifacts, such as tissue folds, antibody aggregates, and particulate contamination, that can confound downstream quantitative analyses. These artifacts manifest as localized aberrant signal intensities, often exacerbated by channel crosstalk (e.g., FITC/Cy3 bleed-through^55^), which amplifies fluorescence overlap and generates spurious bright regions (**Figure 1D**, cancer metastasis in liver example). To mitigate these effects, SpatioEv incorporates a multi-step artifact quality control (QC) module. First, intensity-based thresholding and phenotyping isolate artifact-affected regions based on user-defined signal thresholds (**Figure 1E, Supplemental Figure 1B**). Next, Density-Based Spatial Clustering of Applications with Noise (DBSCAN)^56,57^ identifies and excludes local artifact clusters, preventing skewed cell density measurements due to contamination (**Figure 1F**). This systematic approach ensures artifact-free regions are retained for downstream spatial analyses while preserving data integrity.

Accurate and consistent cell phenotyping across large-scale multiplexed imaging datasets is critical for robust biological interpretation. Technical variations in staining intensity between imaging runs can introduce significant variability in marker expression levels, complicating the identification of distinct cell populations. While manual cell gating is often considered a gold-standard, its scalability is limited, and subjective biases may arise in large datasets. To address these challenges, SpatioEv integrates an automated, unbiased normalization and phenotyping module using Gaussian Mixture Models (GMMs)^58^ to objectively determine marker-negative populations and establish expression thresholds (**Supplemental Figure 1C**). This approach is compatible with standard tools like SCIMAP^35^ for data rescaling and phenotype refinement.

We validated this GMM-based method on a 9-plex B cell imaging dataset across peri-TLS region within pancreatic ductal adenocarcinoma (PDAC) sample comprising 158 fields of view (FOVs), which exhibited pronounced batch effects both within (across FOVs) and between samples. By applying GMM-derived weights to transform the data (**Supplementary Figure 1C**), we effectively mitigated these artifacts. For example, a multiplexed image (**Figure 1G**) using the standard SCIMAP annotation approach, proliferating epithelial cells were markedly overrepresented (**Figure 1H**), whereas GMM-normalized data yielded biologically plausible cell distributions (**Figures 1I**). This demonstrates how unaddressed technical variability can skew cell-type proportions and lead to erroneous interpretations, underscoring the necessity of our high-confidence labelling approach for reliable spatial analyses.

### Enhanced annotation of cell annotations and transition states via semi-supervised SVM

Accurate phenotyping is a crucial prerequisite for robust single-cell spatial analysis. However, inherent challenges such as errors in cell annotation often arise due to signal spillover between neighboring cells. To address this limitation, we developed an approach that integrates SCIMAP^35^ phenotyping workflow with a support vector machine (SVM)^59,60^-based probability model to enhance annotation accuracy and capture cells in transition states (**Figure 2A**). Importantly, this approach uses both marker expression profiles as well as morphological features which are known to be distinct between cell types.

**Figure 2.**
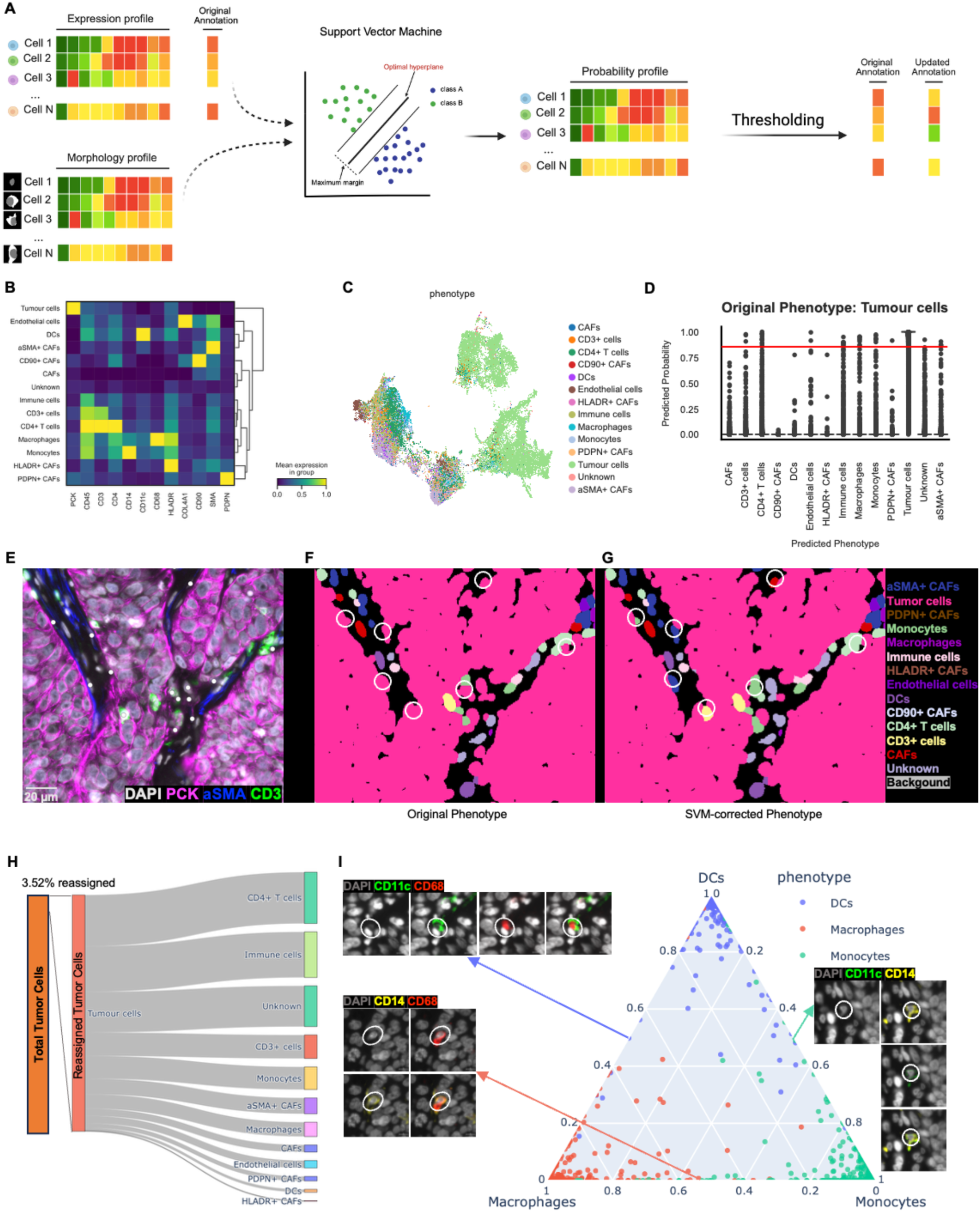
SpatioEv generates high-confidence cell annotations using an SVM-based approach for head and neck cancer imaging data. **A.** Schematic outlining the key steps of the SVM-based method. Expression and morphological profiles are input to generate a probability profile, which is then thresholded to refine cell annotations. **B**. Heatmap illustrating protein signatures for identified cell types. **C**. UMAP visualization of cells colored by their determined cellular phenotype. **D.** A boxplot showing the distribution of predicted probabilities of belonging to each phenotype for tumor cells (original phenotype). **E**. A field of view (FOV) example highlighting tumor cells (original phenotype) with a misassigned phenotype. **F-G**. Corresponding segmentation masks colored by phenotype showing the original (**F**) and updated (**G**) phenotype assignments. **H**. Sankey plot depicting the changes in tumor cells phenotype assignments before and after applying the SVM-based method (only corrected tumor cells are shown here). **I**. Scatter plot showing the probability distribution of Macrophages, Monocytes, and Dendritic cells.

Using a 24-plex head and neck cancer imaging data (acquired through Cell DIVE image system, ∼5.2 x 10^5^ total cells), we preprocessed the data by stitching and merging with ASHLAR^61^, segmenting with Mesmer^53^, and quantified with Pixie^62^. Following SCIMAP-based phenotyping, 14 distinct cell populations were identified (**Figure 2B-C**). To train a semi-supervised SVM classifier, we randomly selected 50% of the dataset as a training set, achieving 92.95% prediction accuracy (**Supplementary Figure 2A-C**). This trained model was then applied to the entire dataset generating phenotype probability scores for each cell (**Supplementary Figure 2D**). To evaluate the model, tumor cell (defined as PCK+ cells) outliers were defined as cells with a tumor phenotype probability score below 0.9 (**Figure 2D**). The model revealed tumor cell (PCK+) misclassifications, defined as cells with low tumor probability scores (<0.9) that were positive CD3 or αSMA indicating that they should be T cells or fibroblasts (**Figure 2E, white dots**). The SVM approach successfully corrected these errors (**Figure 2F-G**). The probability model enabled flexible phenotype refinement: the new phenotype would be defined as the phenotype with the largest probability other than original tumor cells (**Figure 2F-H**).

Beyond cell annotations, the SVM probability model is also effective in identifying cells in transitional states, often missed by traditional approaches^4545^. For example, immune cells and fibroblasts often differentiate or transition between subpopulations in response to microenvironmental cues, a process that categorical phenotyping strategies fail to capture^1^. In the head and neck cancer image dataset, we identified three myeloid populations— macrophages (CD68+), monocytes (CD14+), and dendritic cells (CD11c+). The SVM model calculated phenotype probabilities for each cell (**Supplementary Figure 2D**), revealing some cells with intermediate probabilities, indicating cells potentially undergoing transitions. The resulting distribution plot of the probability scores reveals cells undergoing transitions between cell states (**Figure 2I**), providing a nuanced understanding of cell states and offering a powerful tool for investigating microenvironmental factors associated with cellular transitions.

### Multi-scale spatial characterization of cell distributions within tissue microenvironments

Traditional analyses of cellular microenvironments often rely on cell fractions within defined regions, but this approach fails to capture essential spatial features such as local density variations and inter-population correlations, which subsequently can distort correlations between population distributions. To overcome these limitations, we developed a flexible spatial analysis framework that quantifies cell distribution patterns at multiple scales.

Here, we define three density metrics which can be applied to all cells as well as individual cell types (**Figure 3A**, left). To assess global cellular density, we partitioned images into small regions (default: 128×128 pixel tiles) and computed two complementary density metrics: (1) cell counts per tile (**Figure 3B**) and (2) pixel area coverage, accounting for cell size heterogeneity (**Figure 3C**), demonstrated on 25-plex rheumatoid arthritis (RA) images (**Supplementary Figure 3A**). This dual approach enables phenotype-specific density mapping, as demonstrated for B cells revealing B cell-enriched clusters (**Figure 3D, Supplementary Figure 3B**). This also facilitates correlations between densities of different phenotypes, offering further insights into spatial relationships within the microenvironment, for instance, the positive association between CD4+ T cell and B cell densities (**Figures 3E, Supplementary Figure 3C**). To examine the local spatial of cells at a finer scale, we also developed a local density metric of individual cells (**Figure 3A**, right) which corresponds to the inverse of the mean distance to K-nearest neighbors, applicable to all cells (**Figure 3F**) or specific subsets (**Figure 3G**). This high-resolution approach identifies micro-niches, such as tertiary lymphoid structures, and captures boundary dynamics, such as tumor-stroma borders offering insights inaccessible to fractional abundance methods alone. Although many of these metrics are highly correlated, each provides unique information about tissue organization, enabling discrimination between distinct architectural patterns and underlying biological processes.

**Figure 3.**
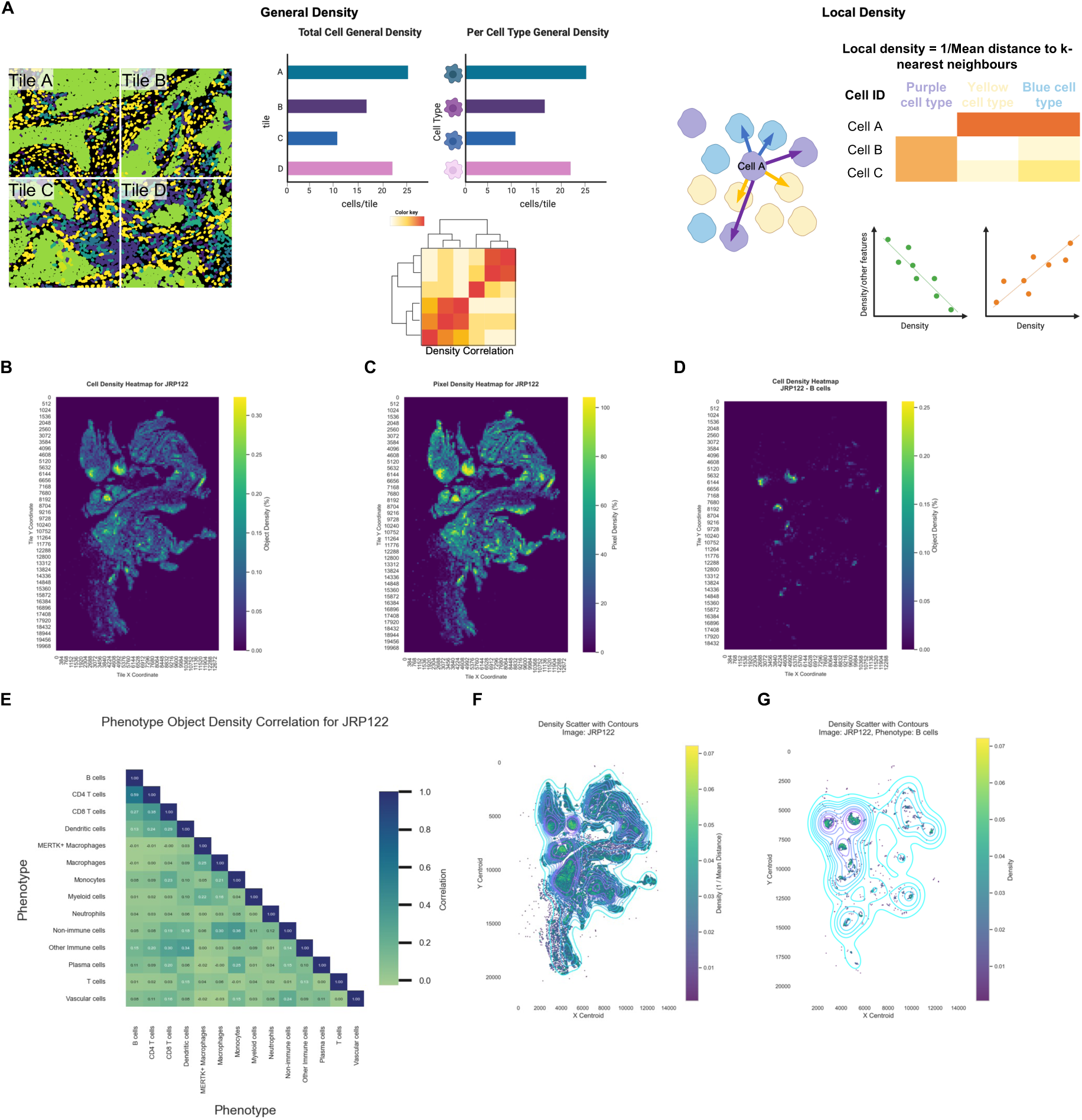
General and local cell density analysis in rheumatoid arthritis (RA) images. **A**. Schematic defining general cell density (tile-based) and local cell density (cell-based) with potential applications. **B**. Representative RA image colored by the general density of all cells (cell counts per tile) (yellow indicates higher density). **C**. The same RA image colored by pixel density (yellow indicates higher density). **D**. RA image colored by the general density of B cells. **E**. Pearson correlation matrix of general cell density across different cell phenotypes. **F**. Scatter plot showing the local density of cells in the RA image, with density contours overlaid. **G**. Density scatter plot specifically for B cells in the RA image, with density contours.

### Automated neighbourhood and region-of-interest selection

Tissue architecture is inherently heterogeneous, composed of distinct cellular neighbourhoods such as immune-rich tertiary lymphoid structures (TLS) or tumour cell-rich tumour nests. Current analytical approaches often depend on manual histopathologist annotations, which introduce subjectivity, inconsistency, and scalability challenges for large datasets. Emerging computational tools, including Giotto^63^, BayesSpace^64^, and Spatial-LDA^65^, address this by algorithmically decomposing spatial imaging data into biologically meaningful regions. SpatioEv enhances this analytical framework by combining Spatial-LDA’s latent Dirichlet allocation topic modelling (which identifies spatially coherent patterns from cell-type distributions) with k-means clustering^66^ and DBSCAN^57^ to standardize neighbourhood definitions across multiple images and connect same neighbourhoods located next to each other. Applied to a cohort of RA and OA with general cell phenotypes annotated (**Supplementary Figure 3A**) and used as input for the neighbourhood identification, this approach identified 10 distinct neighbourhood types (**Figure 4A**) that showed clear spatial segregation (**Figure 4B**) and differential abundance between disease groups (**Supplementary Figure 4B**), enabling automated detection of clinically relevant tissue domains (**Figure 4C**). Tertiary lymphoid structures (TLS), which form in chronically inflamed tissues and resemble secondary lymphoid organs, are characterized by organized aggregates of T cells, B cells, plasma cells, dendritic cells, and specialized stromal components that support local immune activation and memory formation^67^. Within our dataset, several RCNs reflect discrete immune-enriched microenvironments consistent with TLS components: RCN 4 is enriched for B cells, plasma cells, and T cells; RCN 2 for dendritic cells and T cells; RCN 5 for plasma cells; and RCN 6 for T cells, monocytes, and other immune subsets. The spatial co-localization of these RCNs suggests that our approach captures distinct substructures of TLS within synovial tissue, highlighting the fine-grained cellular organization underlying local immune responses in inflamed joints.

**Figure 4.**
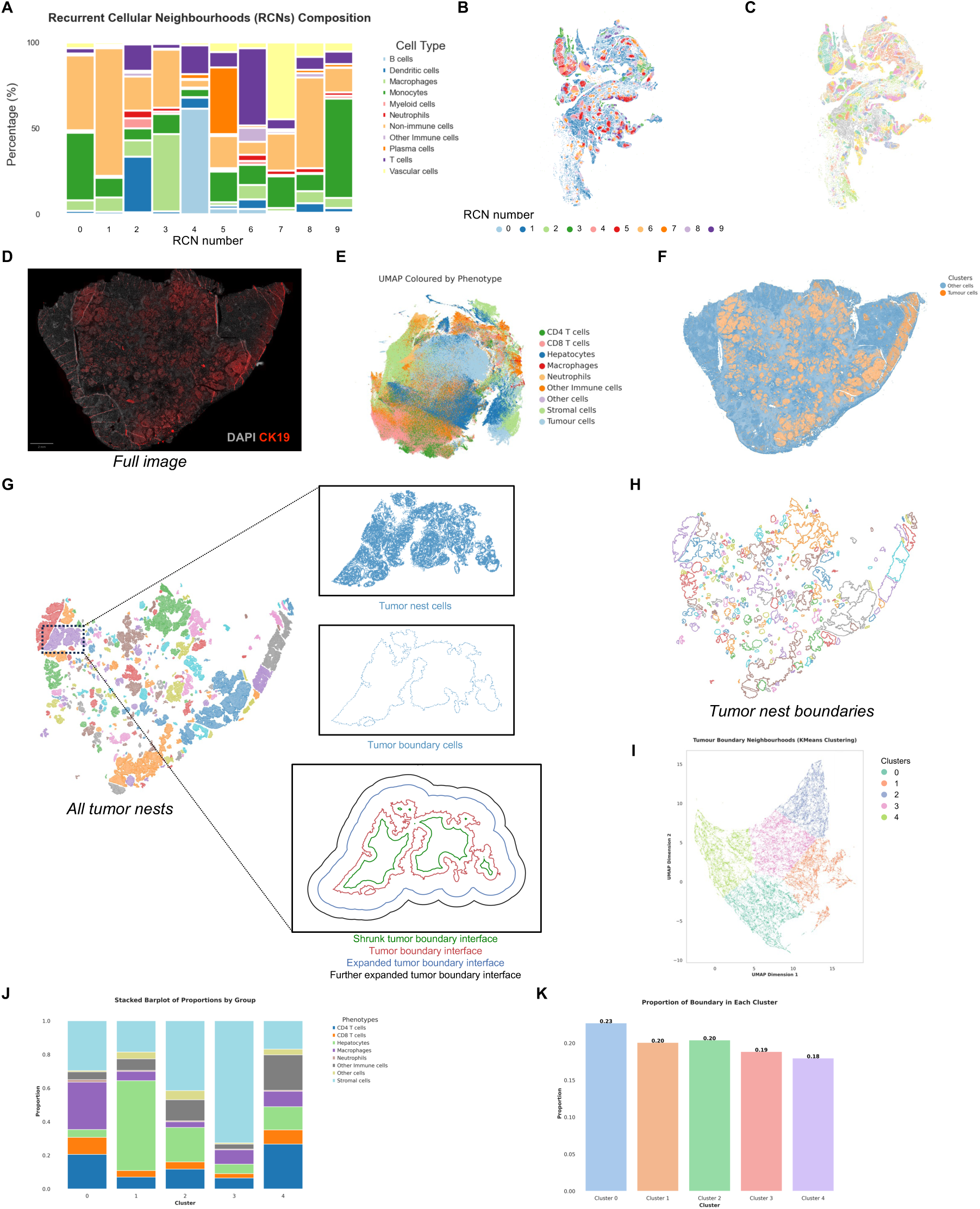
Identification and characterization of tumor nests in cancer metastasis in liver. **A.** Stacked bar plot showing the cell type composition of recurrent cellular neighborhoods (RCNs). **B.** spatial scatter plot colored by RCNs. **C.** Spatial scatter plot colored by connected ROIs. **D.** Representative cancer metastasis in liver image stained with DAPI (nuclei) and CK19 (cytokeratin 19, highlighting ductal cells). **E.** UMAP visualization of cells from the cancer metastasis in liver colored by their identified phenotypes. **F**. An overlay image highlighting clusters of tumor cells and other cells. **G.** Tumor nests defined using DBSCAN based on local cell density and proximity. Zoomed-in views show an individual tumor nest (upper), the defined boundary (middle), and examples of shrunken and expanded boundaries in green and red, respectively (lower). **H.** Tumor nest boundaries across the entire liver metastasis image. **I.** UMAP embedding of boundary cell neighborhoods. **J.** Stacked bar plot illustrating the cellular composition of the boundary cell neighborhoods. **K.** Proportions of cells in different boundary clusters.

### Neighbourhood boundary and interface analysis

The boundaries between cell clusters or neighbourhoods represent critical zones of potential cellular signalling perturbations and interactions^36,39^. For example, the tumour microenvironment (TME) is spatially heterogeneous, with critical interactions occurring between tumour cells and stromal cells at the interface of the tumour nests^1,68,69^. Indeed, these tumour-boundary interactions have been shown to be associated with prognosis in liver metastases^70–75^. Here, we introduce a novel and generalisable module that automates the detection of neighbourhood boundaries enabling interface analysis. Briefly, this approach involves automated neighbourhood annotation (Spatial-LDA^65^ with k-means^66^), DBSCAN-based^57^ clustering of each neighbourhood type to identify local neighbourhood clusters, and boundary identification (via ConcaveHull (shapely.concave_hull v2.0.6)) and boundary quantification.

We applied this analytical framework to a cancer metastasis in liver dataset comprising ∼2 million cells annotated through 24-plex imaging (**Figure 4D**). Current pathological classification of liver metastases relies on binary tumour boundary definitions, typically distinguishing desmoplastic (fibro-inflammatory stroma-demarcated) from replacement (tumour-infiltrated) growth patterns^76^, with the former associated with superior clinical outcomes^72,77,78^. However, these classifications represent extremes of a biological continuum, and the spectrum of tumour-stroma interfaces likely reflects dynamic cellular and molecular interactions that remain poorly characterized at spatial resolution.

Through SCIMAP-based phenotyping, broad cell types were identified (**Figure 4E**, **Supplementary Figure 4A)**. Here, we aimed to understand the heterogeneity of tumour-stromal boundaries. Thus, cells were broadly categorized as either “tumour cells” or “other cells” (**Figure 4F**). Tumour cells were then grouped into spatial clusters based on their physical proximity (via DBSCAN^56,57^), with each cluster of tumour cells representing a distinct tumour nest (**Figure 4G**, left). To delineate tumour boundaries, we applied the ConcaveHull function (shapely.concave_hull v2.0.6), enabling precise identification of edge-located cells within each neighbourhood cluster. Using this boundary framework, we mapped regions that either expanded beyond or contracted within the tumour boundary contour (**Figure 4G** right; **Figure 4H**). SpatioEv allows for customizable parameter tuning, empowering users to refine tumour clustering and boundary segmentation to suit specific biological questions. Finally, to interrogate the spatial context of boundary regions, we characterized the local neighbourhood composition of boundary cells. Leveraging SCIMAP’s spatial_count function, we computed the surrounding cell types after renormalizing the data to exclude tumour cells themselves. This allowed us to focus on the microenvironmental context. The resulting neighbourhood boundary profiles were dimensionally reduced using UMAP^79^, clustered using K-means^66^ (**Figure 4I-J**), and their relative proportions quantified (**Figure 4K**). We identified five distinct tumour boundary types, each characterized by unique cellular neighbourhood compositions (**Figure 4J**). The most prevalent boundary (Cluster 0) consisted predominantly of CD4+ T cells, macrophages, and stromal cells, suggesting active immune-stromal interactions. Notably, Cluster 1 boundaries showed hepatocyte enrichment, indicating direct tumour-parenchyma interfaces, while other clusters exhibited immune-dominant, fibroblast-rich, or mixed cellular profiles. This spectrum of boundary architectures implies varying mechanisms of tumour containment, invasion, and stromal remodelling. Importantly, such boundary heterogeneity may drive differential clinical outcomes, not just between patients, but even among distinct tumour nests within individual patients, potentially influencing both prognosis and therapy response.

This method is adaptable and can be applied easily to other datasets to analysis and other neighborhood types. For example, this can be adapted to identify other structural boundaries’ such as TLSs. This demonstrates SpatioEv’s versatility for exploring spatial boundaries across various tissue types and disease contexts.

### Integrated spatial analysis of extracellular matrix fiber proteins and cellular features

The ECM is increasingly recognized as a dynamic and critical component of tissue microenvironments, exerting profound influences on both physiological homeostasis and disease progression^4,6,28^. Beyond its biochemical composition, the physical architecture of the ECM, including fiber alignment, length, density, and the specific types of constituent proteins, plays a significant role in regulating cellular behaviours such as support, migration, differentiation, and survival^80–82^ . To dissect the intricate spatial relationships between cells and ECM fibers within highly multiplexed imaging datasets, we have developed a novel per-fiber and per-cell analytical module integrated within SpatioEv (**Figure 5A**). This unique approach enables a granular, fiber protein-resolved investigation into how ECM fiber characteristics influence immune cell distribution within the tumour microenvironment (TME) and the underlying mechanisms of tumour cell migration and invasion.

**Figure 5.**
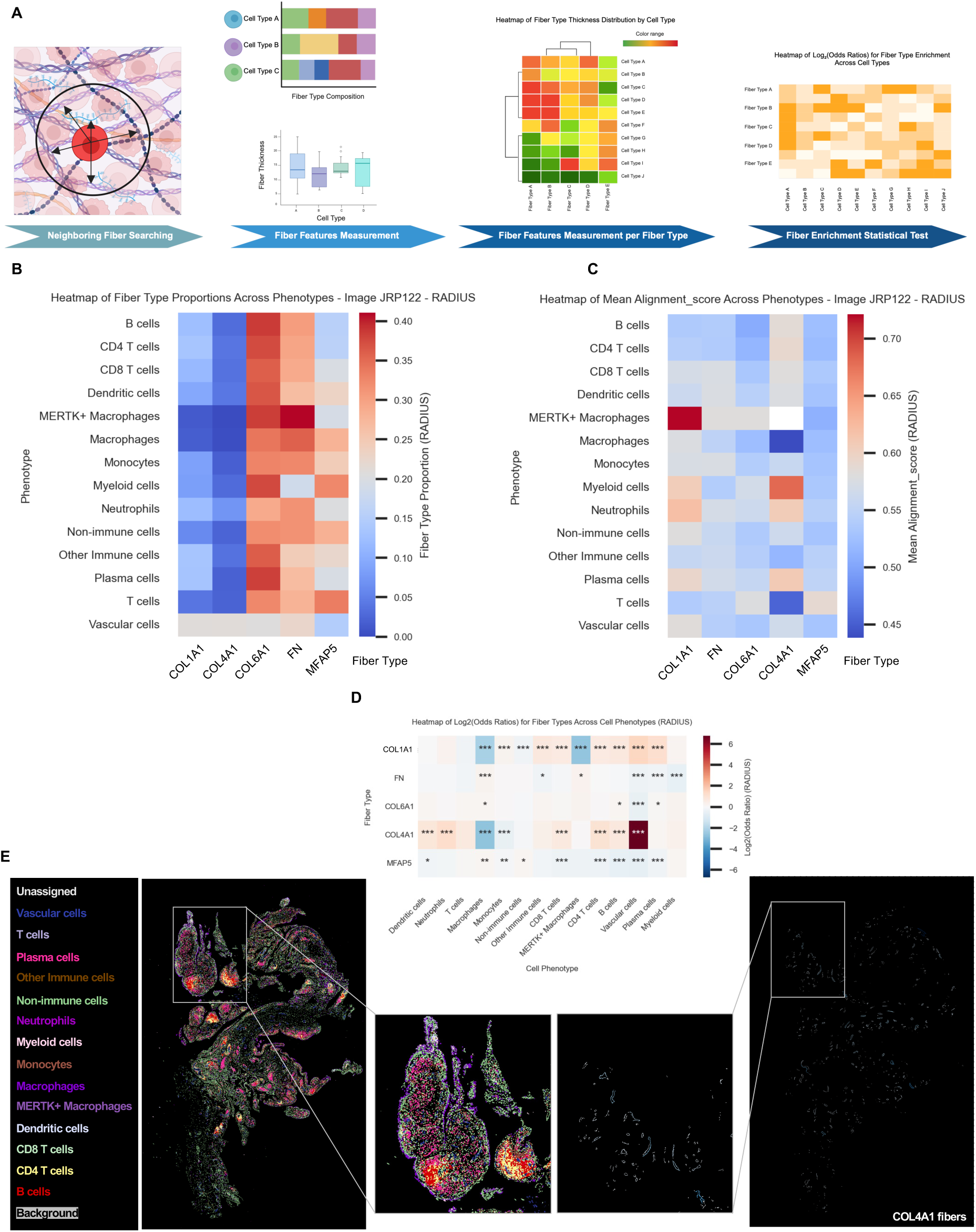
Integrated spatial analysis of extracellular matrix fibers and cellular microenvironments. **A**. Schematic overview illustrating the key analytical steps of the SpatioEv ECM-Cell combined analysis, enabling per-fiber and per-cell spatial investigations. **B**. Heatmap of fiber (COL4A1, COL6A1, MFAP5, FN, COL1A1, CHP) type proportions across phenotypes. **C**. Heatmap displaying mean alignment score across phenotypes. **D**. Heatmap displaying Log2Odds ratio for fiber type across cell phenotypes. **E**. (left) Representative segmentation mask with cells colored by phenotype, with (middle) the spatial localization of B cells (red) in a region of interest, and (right) spatial localization of COL4A1 in the same region of interest.

We applied it to a dataset of three rheumatoid arthritis (RA) and three osteoarthritis (OA) 27-plex images, from which approximately 31,000 high-quality ECM fibers stained with five key ECM markers (COL1A1, COL4A1, COL6A1, Fibronectin (FN), and MFAP5) were segmented using Pixie^62^ and QCed (see *Materials and Methods*). This allows us to identify elongated and segmented structures of fibrillar proteins, which act as a proxy to ECM structure. This allows for the quantification of a morphological and physical features for each fibrillar signal, including area, major axis length, minor axis length, eccentricity, Euler number, and alignment score. UMAP of fibrillar protein segment physical features revealed no significant global differences in these physical features across the different fibrillar protein types (**Supplemental Figures 5A-B**, underscoring the need for spatially resolved analyses.

Next, the proportions of different fibrillar protein types in the immediate vicinity of each cell population can be quantified. SpatioEv module incorporates a flexible and efficient BallTree-based method (sklearn.neighbors.BallTree) for the precise identification of neighbouring fibers for each cell, offering users the choice between k-nearest neighbours^83^ (KNN, k = 5 in example) and radius-based approaches (r = 15 µm in example) (**Figure 5A**). This user-centric design allows for tailored exploration of cell-ECM interactions based on biological context. This reveals how different fiber types and properties (area, major axis length, minor axis length, and alignment score) associate with different cell types (**Figure 5B-C, Supplementary Figure 5C-E**).

To statistically account for different proportions of cell and fibrillar protein segments, our module incorporates a statistical framework based on Fisher’s exact test to calculate enrichment odds ratios, thereby identifying fiber types that are preferentially associated with specific cell populations. This revealed fiber-specific spatial patterning: COL4A1 fibers showed both co-localisation and significant enrichment around vascular cells (OR=107.48, p<0.001; **Figures 5D-E**), consistent with their known vascular basement membrane role. Notably, we identified previously unrecognized associations, including a higher alignment of COL1A1 fibers near MERTK+ macrophages and other myeloid cells (**Figure 5D**) and longer COL6A1 fibers near vascular cells (**Figure 5D**). Our analysis also revealed a significant enrichment of COL4A1 fibers in the vicinity of B cells (p < 0.001, Fisher’s exact test) (**Figures 5D-E**). SpatioEv’s granular spatial analysis capabilities suggests a previously unappreciated interaction between COL4A1-rich ECM structures and B cells within RA. This capability provides a powerful lens for uncovering novel spatial enrichment patterns of ECM fibers and their reciprocal influence on diverse cell populations within complex tissue microenvironments and provides new opportunities to develop stroma-targeted therapies.

### Spatiotemporal trajectory inference of recurrent spatial niches

Given the recurrence of similar tissue niches (‘neighbourhoods’) across whole-slide images, each at varying maturation stages and exhibiting cellular and structural heterogeneity, we developed a novel spatiotemporal trajectory inference framework to model their dynamic evolution and uncover how these niches emerge and progress over time. For example, unique advantage of WSI of tumour sections is its capacity to capture a spectrum of tumour progression stages or TLS maturation states within a single tissue section, offering an unprecedented opportunity to reconstruct the developmental trajectory of cancer at the tissue architecture level^84^. Current understanding of tumour progression largely stems from cross-sectional studies comparing samples between individuals or from animal models, which is limited by inter-patient variability and reduced statistical power^85^. To overcome these constraints, we developed a novel analytical framework that reconstructs tumour evolution within individual patients, while simultaneously characterizing dynamic interactions with the extracellular matrix and microenvironmental cell populations.

To systematically dissect the coordinated evolution of malignant niches and their surrounding microenvironments, we developed an analytical module that: (1) extracts individual tumour nests, (2) quantifies single-cell morphological and molecular features, and (3) characterizes both nest architecture and surrounding cellular context (**Figure 6A**). This integrated approach enables systematic investigation of tumour-stroma dynamics at multiple spatial scales. Thus, we applied this to a 24-plex pancreatic WSI from a pancreatic ductal adenocarcinoma (PDAC) patient, an ideal model system with clearly defined progression from non-invasive pancreatic intraepithelial neoplasia (PanIN) lesions, graded 1-3 by increasing cytological and architectural atypia, to invasive carcinoma. This allows for direct mapping of spatial niche development across tumorigenesis. This stepwise morphological transformation underscores the critical role of tissue-level alterations in disease evolution, a concept central to the tissue organization field theory of oncogenesis, which posits that aberrant stromal-tumour cell interactions drive cancer development. Notably, distinct subtypes of cancer-associated fibroblasts (CAFs) have been shown to emerge early in PanIN pathogenesis^86–88^, highlighting the importance of analysing the evolving cellular ecosystem.

**Figure 6.**
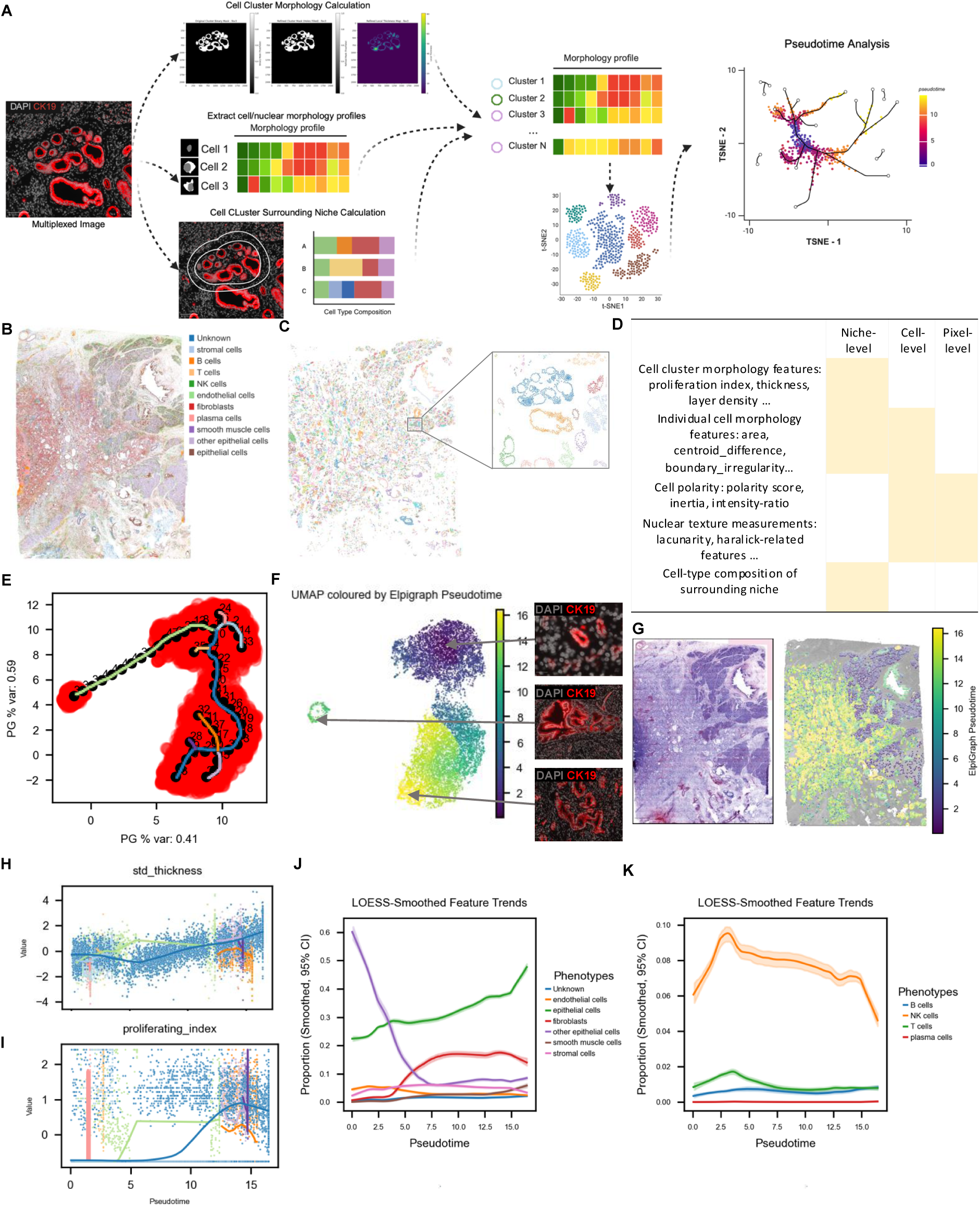
Spatiotemporal Trajectory Inference Reveals Divergent Evolutionary Pathways and Dynamic Niche Composition in Pancreatic Ductal Adenocarcinoma. **A.** Schematic outlining the workflow of our niche-level spatiotemporal trajectory inference analysis, integrating morphological features and microenvironmental context. **B.** Spatial distribution of annotated cell phenotypes within the PDAC WSI displayed in a scatterplot. **C.** Identification of distinct epithelial clusters across the tissue section, with each color representing an independent cluster. **D.** Schematic table of niche-, cellular-, and nuclear-level parameters used for the Spatiotemporal Trajectory Inference. **E**. UMAP plot of epithelial clusters, with the overlaid ElPiGraph-inferred developmental trajectory, illustrating potential branching pathways (by color). **F**. UMAP of epithelial clusters colored by pseudotime values along the trajectory (left); representative images of epithelial clusters with low (right upper), middle (right middle), high (right bottom) pseudotime value. **G**. Virtual H&E image (left) and scatter plot of epithelial clusters colored by pseudotime value (right). **H**. Trends of cluster-level parameters standard deviation of thickness plotted over pseudotime. **I.** Trends of cluster-level parameters proliferating index plotted over pseudotime. **J.** Trends in the proportion of key surrounding non-immune cell types including endothelial cells, epithelial cells, fibroblasts, other epithelial cells, smooth muscle cells, and stroma cells around epithelial clusters along the pseudotime trajectory. **K.** Trends in the proportion of key surrounding immune cell types including T cells, B cells, NK cells, and plasma cells around epithelial clusters along the pseudotime trajectory.

Eleven major cell populations were annotated, encompassing both adjacent normal pancreas and invasive cancerous regions (**Figure 6B, Supplemental Figure 6A**). Pancreatic epithelial cells (including tumour cells and non-tumour cells) were then grouped into spatial clusters based on their physical proximity (via DBSCAN) to define discrete epithelial (or tumour) niches (**Figure 6C**). For each of these niches, we comprehensively quantified a range of niche-, cell-, and nuclear-level parameters (**Figure 6D, Supplementary Table 6**), including thickness, layer density, cell centroid difference, nuclear-to-cell ratio, and circularity capturing pixel-level changes in the arrangement of DNA within the nucleus, which are a hallmark of malignant transformation^89,90^. Cell-level and nuclear-level features were aggregated at the niche level via mean and standard deviation (**Supplementary Table 6**). Crucially, our method incorporates the surrounding cell type composition for each epithelial cell cluster, capturing the dynamic interplay within the tumour microenvironment.

Following normalization and transformation of this integrated feature matrix, we employed UMAP for dimensionality reduction and ElPiGraph^91^ to infer the developmental trajectory of these epithelial niches (**Supplementary Figure 6B-C**, **Figure 6E**). Notably, tumorigenesis has been shown to not follow a linear path but can follow multiple branching pathways^92–95^, a complexity captured by our model (**Figure 6E**). Projecting the inferred pseudotime values onto the UMAP embedding revealed a clear progression, correlating with changes in niche thickness and proliferation along the trajectory, in line with classical histological classifications (**Figure 6F left**, **Supplementary Figure 6D-F**). Finally, projecting the inferred pseudotime values onto the original location of the epithelial clusters clearly showed that normal ductal clusters have the lowest pseudotime values within the adjacent-normal regions of the tissue, aligning with histological annotations (**Figure 6G**).

Next, we explored which features correlated with increased epithelial cluster pseudotime. Quantitative analysis of key cluster-level morphological parameters demonstrated a significant increase in mean and standard deviation of thickness, and proliferation index along the pseudotime axis (**Figure 6H-I, Supplementary Figure 6G-I**), indicating a progressive increase in proliferation, and overall size of epithelial niches, consistent with the established pathological hallmarks of PDAC progression.

At the single-cell level within these niches, we observed increasing trends in parameters such as the mean and standard deviation of cell area, centroid-difference (a measure of cell polarity), equivalent diameter, major axis diameter ratio, perimeter square over area, boundary irregularity, and nuclear-to-cytoplasmic ratio (**Supplementary Figure 7A-G**), signifying a growing morphological heterogeneity, structural irregularity and disorganization of individual epithelial cells as the tumour progresses. Furthermore, the composition of the surrounding cellular microenvironment exhibited dynamic shifts along the inferred pseudotime trajectory (**Figure 6J-K, Supplementary Figure 8A-I**), including the reduction of non-tumour epithelial cells (CK19-CD11b+) and increase in fibroblasts with tumour development. The nature of the fibroblasts varies with tumour niche development (**Supplementary Figure 8C-F**), with a significant reduction in the expression of aSMA but increased expression of Thy1, FAP and PDPN, which are associated change in fibroblast pro-tumorigenesis function^96–100^. Of the immune cells, NK cell proximity to tumour clusters increases at early pseudotime but decreases significantly with more progressed tumour nests suggesting a switch from immunogenic to immunosuppressive niches. CD4:CD8 T cell ratios change with tumour progression favouring CD4 T cells in more advanced tumour niches (**Supplemental Figures 8A-B**)^101^. Finally, B cells have a higher expression of cytotoxic (GZMB) and immunosuppressive (PDL1) markers^102,103^ (**Supplemental Figures 8G-H**). Together this directly supports the notion of evolving reciprocal interactions between epithelial cells and their niche during tumour progression.

We also identified morphological-level alterations associated with tumour progression. The cell polarity score, defined as the normalized distance between geometric and intensity-weighted centroids in the CK19 channel, captures disruption of apical–basal polarity, a hallmark of epithelial transformation (**Supplementary Figure 9A**) shows a significant decrease with tumorigenesis. The moment of inertia measurement differentiates between compact and diffuse patterns of protein localization (**Supplementary Figure 9B**), revealing a shift toward more diffuse localization during tumorigenesis. Co-localisation metrics intensity ratios between CK19 and NaKATPase revealed an early and rapid loss of membrane marker co-localisation distribution (**Supplementary Figure 9C-D**).

Nuclear texture features derived from the DAPI channel, such as Shannon entropy, lacunarity, and Haralick metrics, quantify chromatin organization, with higher values indicating greater heterogeneity often linked to transcriptional activation or neoplastic transformation (**Supplementary Figure 9E**–J). When projected along the pseudotime trajectory, these features revealed consistent patterns of chromatin remodeling, polarity loss, and spatial disorganization, aligning with known pathological transitions^104,105^ and exposing subtle phenotypic changes not detectable through cell type–based analyses. These changes arise from altered mechanical forces, defects in nuclear structure (e.g. the lamina), and disrupted chromatin organization.

Our spatiotemporal trajectory inference method provides a powerful and quantitative framework to map tumour progression at the niche level, integrating epithelial morphology, dynamic microenvironmental composition, and detailed cellular and subcellular features. This comprehensive approach enables the characterization of multiple, potentially divergent tumour evolution pathways, highlighting the inherent complexity and heterogeneity of tumorigenesis and offering a unique lens for understanding its multifaceted progression.

### Unsupervised weighted feature correlated network analysis (WFCNA) for identifying spatial signatures between image groups

To enable interpretable and statistical comparisons of spatial imaging data across groups, we developed an unsupervised, multi-scale framework for feature extraction and statistical framework. This pipeline encodes a broad array of spatial features, capturing the composition, morphology, and spatial relationships of both cellular and acellular components across hierarchical levels, including individual cells, ECM, cell neighbourhoods, tissue boundaries, and overall tissue architecture to generate a high-dimensional ‘spatial architecture map’ (SAM) encapsulating diverse spatial characteristics of each image (**Figure 7A**, left).

**Figure 7.**
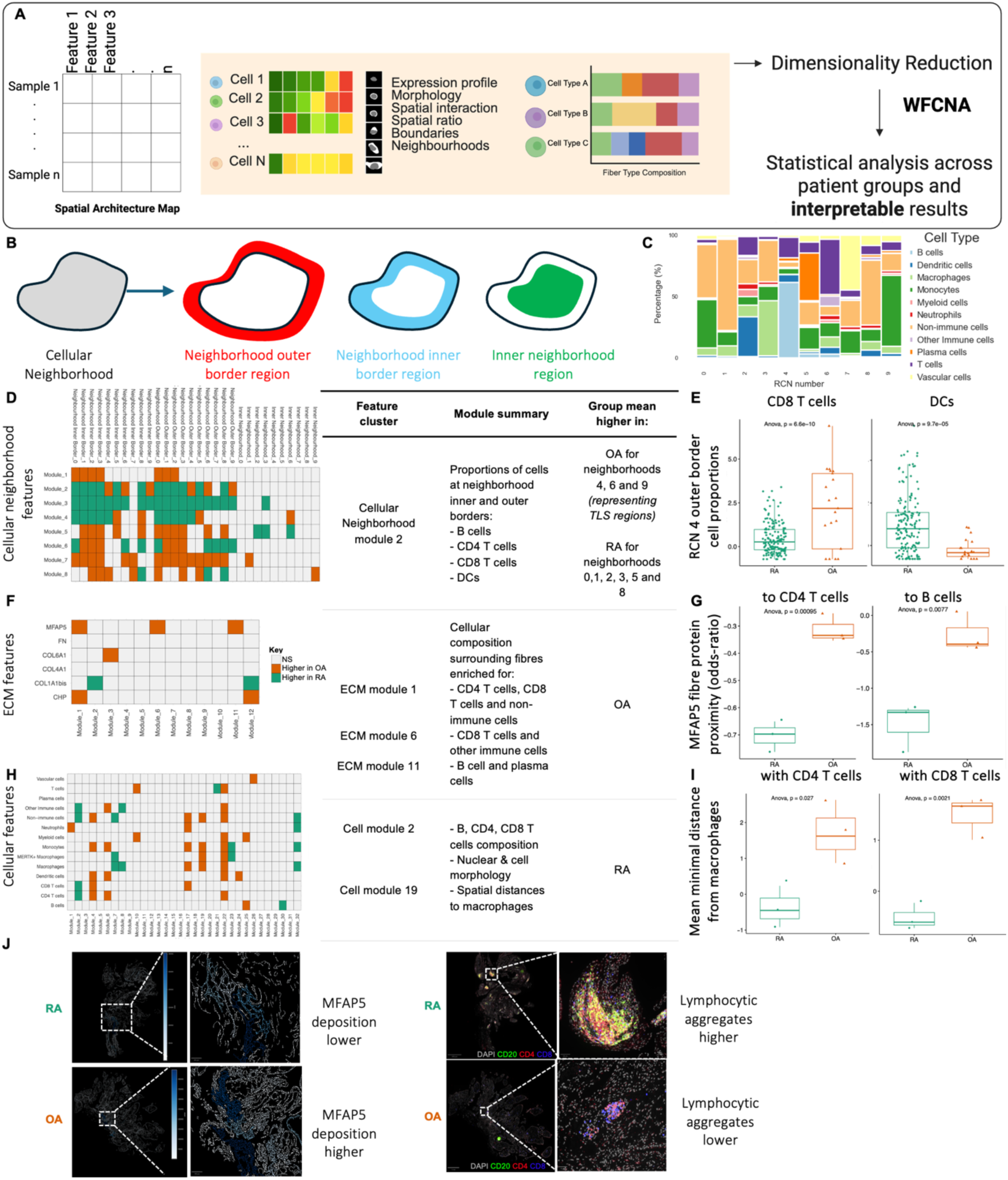
Weighted Feature Co-expression Network Analysis (WFCNA) Reveals Spatial Feature Modules Associated with Rheumatoid and Osteoarthritis. **A.** Schematic of the WFCNA pipeline. A multi-scale spatial feature matrix (Spatial Architecture Map) is generated from images and used as input. Dimensionality reduction via WFCNA groups highly correlated features into distinct modules, which are then subjected to statistical analysis to derive interpretable biological insights. **B.** Schematic of definition of cellular neighborhood regions used in the WFCNA. **C.** Stacked bar plot showing the cell type composition of recurrent cellular neighborhoods (RCNs) in the RA and OA image dataset. Identification and characterization of WFCNA modules at different biological scales (left), example module meaning (middle) and example distribution of selected module features (right) for **D-E.** cellular neighborhood-level features, **F-G.** ECM-level features and **H-I**. cellular-level features. Left: Heatmaps illustrate feature (rows) membership within identified WFCNA modules (columns). Center: For a selected representative module (e.g., Module 3 from ECM features enriched in OA; Module 2 from cell-type features enriched in RA; Module 2 from neighborhood features enriched in RA; all module significance FDR < 0.05), key contributing features are listed. Right: Boxplots show the distribution for an example feature from a significantly different module, comparing RA (teal) and OA (orange) groups (ANOVA p-values indicated). **J**. Representative immunofluorescence images illustrating ECM (left) and cellular-level and cellular neighborhood-level derived signatures.

However, the resulting feature space consists of tens of thousands of descriptors per image, introducing challenges related to multiple hypothesis testing and interpretability. To address these issues, dimensionality reduction is performed using the principles of Weighted Gene Correlation Network Analysis (WGCNA)^106^ to the spatial domain. This approach groups highly correlated spatial features into distinct modules, each representing a coherent spatial pattern (**Figure 7A**, right). By performing statistical analyses at the module (eigenvector) level, we substantially reduce the data complexity and burden of multiple testing while enabling biological interpretation through analysing the enrichment of feature types within each module.

We applied this approach to the 27-plex images from three RA and three OA patients. The complex pathophysiology manifests not only with altered cell composition but also in disruption and reorganisation of tissue architecture. To quantitatively capture these changes, we employ our novel unsupervised Weighted Feature Correlated Network Analysis (WFCNA) dissecting tissue environment into distinct layers: cellular neighbourhood, cellular, and ECM level (**Figure 7B-I, Supplemental Tables 7-9)**.

To examine broad tissue-level organizational patterns, we applied WFCNA to 22 features summarizing cell and ECM composition within different cellular neighborhood types, as well at the border regions where the neighbourhood interfaces with other neighbourhoods (inner neighbourhood region, neighbourhood inner border, and outer border, **Figure 7B**). This analysis uncovered seven modules, all of which were significantly associated with disease state (**Figure 7C-E**). For example, Module 2 reflects the proportions of B cells, CD4 T cells and CD8 T cells which were elevated in OA for neighborhoods 4, 6 and 9, representing the TLS regions, and RA for neighborhoods 0,1, 2, 3, 5 and 8 representing predominantly non-TLS tissue regions. This therefore reflects coordinated shifts in tissue architecture, particularly at neighborhood interfaces, indicating a global reorganization of the immune landscape rather than changes limited to a specific region.

To dissect coordinated ECM remodelling, we applied WFCNA to 56 neighbourhood enrichment metrics characterizing the spatial context around major ECM fibers using KNN- and radius-based methods. This revealed 12 spatial modules of which 6 differed significantly between RA and OA with at least one fiber type (**Figure 7F-G, J**). Notably, we uncovered novel disease-specific ECM-immune interactions: firstly, MFAP5 fibers exhibit closer spatial proximity to CD4⁺ and CD8⁺ T cells (modules 1 and 6), as well as to B cells and plasma cells (module 11), suggesting disease-specific immune-ECM interactions. While prior studies have indicated that MFAP5 contributes to fibrosis, elevated IL-6 levels, and cartilage degeneration in mouse models^107,108^, our study is the first to reveal distinct differences in MFAP5 fiber architecture and immune cell positioning between OA and RA. Secondly, we observe increased colocalization of collagen hybridizing peptides (CHPs), which selectively bind to denatured collagen and a hallmark of OA-associated tissue degradation^109,110^, with CD4⁺ and CD8⁺ T cells in OA relative to RA (module 1), highlighting a potential mechanism linking ECM damage to immune cell recruitment and sustained inflammatory responses. These findings expand the growing appreciation of ECM’s immunomodulatory roles^111,112^ and suggests new mechanisms whereby matrix composition may shape lymphoid aggregate formation in autoimmunity^113^.

We finally explored disease-specific shifts in cell-intrinsic features and their spatial context. We extracted summary statistics (mean, median, SD) for marker expression, morphology, and spatial features across defined cell phenotypes. WFCNA identified multiple disease-associated modules (**Figure 7H-I**). For example, cellular module 19 reveals a significantly larger distance between macrophages and CD4 and CD8 T cells in OA. Module 2 was significantly enriched in RA samples (FDR < 0.05) and contained co-varying features of B cell, CD4, CD8 T cell composition, nuclear and cell border morphology complexity, and myeloid, monocyte, CD4 and CD8 T cell spatial ratios. These modules features reflect the formation of dense lymphocytic aggregates, ectopic lymphoid structure, and cellular proliferation, which are hallmarks of structural remodelling typical of inflamed tissues and RA pathology ^114,115^ (**Figure 7J, right**).

Crucially, this network-based approach reveals non-obvious spatial signature, for instance, integrating multiple ECM features into an “OA-fibrosis” module and clustering diverse immune markers into an “RA-inflammation” module (**Figure 7J**). These modular patterns would be difficult to detect through traditional single-marker analyses, highlighting the power of WFCNA to resolve complex, multi-scale tissue remodelling in disease.

## Discussion

Our study introduces SpatioEv, a comprehensive computational framework that addresses critical gaps in spatial analysis of highly multiplexed tissue imaging data. By integrating automated quality control, probabilistic cell phenotyping, automated neighbourhood identification, multi-scale spatial characterization, niche boundary analysis, ECM fiber-cell interaction mapping, and spatiotemporal trajectory inference, SpatioEv enables systematic exploration of tissue organization at unprecedented resolution. Our findings demonstrate how spatial context fundamentally shapes cellular behaviour and tissue function, reinforcing emerging paradigms in spatial biology while providing novel analytical capabilities to test these concepts empirically. This unified pipeline addresses key analytical challenges in the field and provides a powerful platform for revealing the complex spatial architecture and dynamic cellular interactions that underpin tissue organisation, disease progression and enables a more holistic interrogation of the complex spatial relationships within tissue microenvironments.

Our automated quality control and phenotyping modules address longstanding reproducibility issues in image analysis. By implementing Gaussian mixture models for marker thresholding and SVM-based probability scoring, we were able to rapidly and accurately annotate cell types while capturing transitional cell states, a significant advance over manual gating approaches that frequently miss these biologically important populations. This is particularly relevant for understanding immune cell differentiation and stromal cell plasticity in diseases like rheumatoid arthritis and cancer.

The multi-scale spatial analysis capabilities of SpatioEv revealed several novel insights into tissue organization. Our density mapping approach demonstrated that immune cell interactions in RA tissue occur at characteristic spatial scales, B and T cell correlations emerged when analysing local neighbourhoods (50-100 μm), underscoring the importance of scale-sensitive analysis. This aligns with known B and T cell interactions within TLSs. Perhaps most strikingly, our ECM analysis uncovered previously unrecognized fiber-cell interactions, including COL4A1 enrichment around B cells in RA. This finding expands growing appreciation of ECM’s immunomodulatory roles^111,112^ and suggests new mechanisms whereby matrix composition may shape lymphoid aggregate formation in autoimmunity^113^. The ability to analyse fiber characteristics at single-cell resolution represents a major technical advance over existing ECM analysis methods^116^.

Tissue architecture is highly heterogeneous, and while traditional annotation methods are limited by subjectivity and scalability, SpatioEv builds on advanced computational tools to standardize and automate the identification of biologically meaningful tissue neighbourhoods, enabling reproducible, clinically relevant spatial profiling across large datasets. This advance enables a novel, generalizable module within SpatioEv that automates the detection and characterization of tissue neighbourhood boundaries, which are critical zones of cellular interaction, using spatial clustering and boundary quantification. Applying this to cancer metastases on liver, we identified five distinct tumour boundary types with unique microenvironmental compositions, revealing clinically relevant spatial heterogeneity that may influence prognosis and therapeutic response, and demonstrating broad adaptability across disease contexts. More broadly, SpatioEv’s ability to quantify boundary phenotypes enables precise characterization of tumour-stroma interfaces—a critical determinant of metastatic behaviour^52^ .

Our novel spatiotemporal trajectory framework provides the first method for inferring developmental progression in unprecedented detail directly from spatial imaging data using both morphological and cellular characteristics. Application to PDAC revealed branching evolutionary pathways with distinct stromal remodelling patterns, proliferation, and immune infiltration profiles associated with different progression routes. Furthermore, we were able to disentangle early changes in tumorigenesis (loss of membrane CK19 and NaKATPase co-localisation distribution, CD4:CD8 T cell ratio, NK cell increase, non-tumour epithelial cells (CK19-CD11b+) decrease, nuclear morphology features), intermediate or gradual changes (epithelial cluster niche morphology, size, and irregularity, increase in fibroblasts), and late changes during tumour development (reduction in NK cells, proliferation of epithelial cells. The early changes in tumorigenesis may serve to be sensitive markers for early diagnosis. This supports emerging concepts of parallel tumour evolution while providing spatial context missing from sequencing-based lineage tracing. The observed correlation between niche disorganization and immune exclusion offers new insights into immunotherapy resistance mechanisms. By focusing on within-patient, inter-tumour niche analysis, our approach reduces confounding factors from host genetic variability and immune responses, while benefiting from enhanced statistical power due to the large number of tumour niches (up to ∼1000s per image) captured per patient, which is an advantage difficult to achieve in between-patient comparisons. This framework is scalable and generalizable, with potential for cross-patient trajectory mapping across images to uncover shared and divergent pathways of niche evolution. Moreover, it can be readily adapted to study other recurring spatial niches, such as bacterial colonies or tertiary lymphoid structures (TLSs), including distinctions between immunogenic and immunosuppressive TLS development.

To address the challenge of interpreting high-dimensional spatial imaging data across disease states, we developed an unsupervised, multi-scale framework (WFCNA) that enables statistically robust and interpretable comparisons across whole-slide images. By integrating diverse spatial features at the cellular, neighbourhood, and ECM levels, WFCNA organizes complex spatial information into biologically meaningful modules. This approach significantly enhances interpretability while mitigating the challenges of multiple hypothesis testing. By condensing thousands of spatial descriptors into a small number of disease-relevant modules, WFCNA offers a powerful tool for uncovering novel spatial biomarkers and understanding the coordinated remodelling of tissue architecture in disease and offers new biological insights into disease-specific tissue remodelling. For instance, we reveal MFAP5 fibers in RA show increased proximity to T and B cells, suggesting a role in immune cell localization not previously described. In OA, we find enhanced colocalization of denatured collagen (CHPs) with T cells, linking ECM degradation to immune recruitment and chronic inflammation. These findings reveal new mechanisms by which ECM architecture may shape immune responses in joint disease. We also show coordinated shifts in tissue architecture, particularly at neighborhood interfaces, indicating a global reorganization of the immune landscape rather than changes limited to a specific region. Importantly, WFCNA captured a quantitative and integrated view of global tissue immune reorganization in RA, not limited to local aggregates but spanning entire tissue compartments.

While SpatioEv significantly advances the field of spatial biology, some limitations remain. The computational resources required for analysing large-scale, high-dimensional spatial datasets can be substantial. Computational demands increase with image size, however, many of the steps presented here facilitates parallel processing and the modular design of the pipeline allows for user adaption at each step. Furthermore, the accuracy of downstream analyses is inherently dependent on the fidelity of cell and/or ECM segmentation and the comprehensiveness of the marker panel employed. Whilst we address this with the automated QC modules, image quality and manual inspection to check accuracy is still required. Future integration with spatial transcriptomics could further enhance molecular resolution.

By bridging critical gaps in spatial data analysis, SpatioEv empowers researchers to explore tissue organization with unprecedented breadth and precision. The framework’s modular design and integration with the existing SCIMAP framework ensures adaptability to diverse biological questions while maintaining analytical rigor. As multiplexed imaging becomes increasingly central to biomedical research and clinical decision making, tools like SpatioEv will be essential for unlocking the full potential of spatial biology to understand tissue function and dysfunction.

## Materials and Methods

### SpatioEv framework

The SpatioEv framework is designed for accessibility and is accompanied by step-by-step Jupyter notebooks, publicly available on GitHub. These notebooks facilitate a modular workflow, allowing users to initiate analysis at various stages. Input data is required in the Anndata format. The repository includes specifications for the necessary software environments, and each step details the libraries, versions, and parameters used.

### Sample access and preparation

Multiplexed imaging datasets will be available upon reasonable request.

### Image acquisition and pre-processing

To enable accurate image stitching and registration by ASHLAR, we first generated OME-XML companion files for each imaging cycle from Cell DIVE raw imaging tiles. This step was needed to extract and encode spatial metadata (stage positions and pixel size) not natively interpretable by ASHLAR.

We customized a Python script (multi-ome-xml-single-channel-celldive-2.py) that performs the following operations for each imaging cycle folder: it parses TIFF metadata to extract x and y stage positions from the image headers, constructs OME-compliant metadata structures using the ome-types and tifffile libraries, and programmatically appends physical pixel sizes and per-channel spatial positioning information to a synthetic OME image model. Each OME object is assigned a unique UUID to ensure traceability. The script generates one OME companion file per cycle, consolidating metadata from all channels present (default: DAPI, FITC, Cy3, Cy5), and associates each image tile with its correct physical location and channel index.

To automate processing across multiple imaging cycles, we wrote a Bash wrapper script (ome_companion_generate.sh) that loops through all subdirectories matching the S0* naming pattern which are raw file folders generated from Cell DIVE imaging and invokes the Python script with appropriate arguments. This pipeline was executed in a high-performance computing environment using SLURM (srun) with appropriate module loads (Anaconda3 and Java) and a dedicated Conda environment (ashlar) to ensure reproducibility and compatibility with downstream ASHLAR registration.

The stitched and merged ome.tiff format files are then processed with the background subtraction function (module: background) embedded in MCMICRO with exposure time difference between staining cycle and baseline cycle (background round and bleaching rounds) considered.

Whole-cell and nuclear segmentation are performed with Mesmer embedded in MCMICRO. The markers used for the segmentation are provided in the Dataset description part.

Regarding the quantification of all the markers and morphological features extraction, we adapted the source codes of Pixie by adding ’orientation’, ’solidity’, ’feret_diameter_max’, ’circularity’, ’fractal_dimension’, and ’boundary_irregularity’, in the settings.py and their calculations in regionprops_extraction.py.

### Dataset 1

Head and Neck Cancer - Antibody panel is attached as Supplementary Table 1.

Multiplexed images were acquired with Cell DIVE platform.

#### Slide clearing, antigen retrieval, and blocking

Formalin-fixed paraffin-embedded (FFPE) tissue sections (5 μm thick) were baked at 60 °C overnight. Slides were subsequently deparaffinized using xylene and rehydrated through a graded ethanol series. Permeabilization was performed using 0.05% Triton X-100 in PBS for 15 minutes, followed by PBS washes. Antigen retrieval was carried out in a NxGen Decloaking Chamber (Biocare Medical) using boiling citrate buffer (pH 6, Agilent) and Tris-based buffer (pH 9) for 20 minutes each. Slides were then blocked overnight at 4 °C with a blocking solution containing 1% bovine serum albumin (BSA), 1% donkey serum (Bio-Rad), and 1% goat serum. After washing with PBS, sections were stained with DAPI, washed again in PBS, and coverslipped with mounting media composed of 50% glycerol and 4% propyl gallate (Sigma).

### Background imaging acquisition

The GE Cell DIVE system was employed to acquire the images of FFPE slides. A scan plan was generated at ×10 magnification to define the region-of-interest (ROI), followed by imaging at ×20 magnification to acquire background autofluorescence information and generate virtual H&E images. Background imaging was used to subtract autofluorescence for the first staining round. Slides were de-coverslipped before staining.

### Staining and bleaching imaging acquisition

Slides were first incubated with Fc-blocking solution at room temperature for 1 hour, followed by overnight incubation at 4 °C with a cocktail of three primary antibodies, applied at the concentrations specified in Supplementary Table X. After three washes with PBS, slides were coverslipped with mounting media, and the first round of staining was imaged. Slides were then de-coverslipped and subjected to a bleaching procedure using a solution of 0.1 M sodium bicarbonate (NaHCO₃, pH 11.2, Sigma) and 3% hydrogen peroxide (H₂O₂, Merck), applied twice to quench fluorophores. After three additional PBS washes, slides were stained with DAPI, washed with PBS, and re-coverslipped. Imaging was performed following this bleaching step to acquire an autofluorescence baseline for the subsequent round of staining. Slides were then de-coverslipped again, washed once with PBST (1× PBS containing 0.05% Tween-20) and twice with PBS, and blocked at room temperature for 1 hour using a solution containing 1% BSA, 1% donkey serum (Bio-Rad), and 1% goat serum. Slides were incubated overnight at 4 °C with the next set of three antibodies, as listed in Supplementary Table X. This cycle of staining, bleaching, and imaging was repeated iteratively for subsequent rounds.

### Dataset 2

Pancreatic Ductal Adenocarcinoma - Antibody panel is attached as Supplementary Table 2.

We performed manual mIF staining using the TSA methodology to detect antigens and amplify the signal detection. TSA amplifies immune-fluorescent (IF) Opal polymer horse radish peroxidase (HRP) to enzymatically convert TSA molecules into free radicals that bind to nearby tyrosine sidechain residues, on antigenic determinants targeted by the primary antibody. After labelling is complete, the antibodies can be removed without disrupting the fingerprint Opal IF signal allowing for the next target to be detected without antibody cross-reactivity. Each section was 4μm thick and the slides were baked overnight at 60 °C and then washed in Xylene three times for 10 minutes, followed by ethanol rehydration (Fisher Scientific) and de-ionised water washes. The tissue on the slides were then fixed in 10% neutral buffered formalin (Sigma) for 20 minutes. Akoya Biosciences provides two antigen retrieval buffers, one with pH6 (AR6, Akoya Biosciences) and the other with pH9 (AR9, Akoya Biosciences). The decision on which antigen retrieval buffer to use for each antibody was made by using the same pH of the buffer as used in the chromogenic immunohistochemistry methods. The slides were rinsed in either AR6 or AR9 and then placed in a Hellen dhal staining jar and high-intensity epitope retrieval was performed by boiling the slides in a microwave at full power for one minute. Followed by 15 minutes at low power, ensuring that the evaporation of buffer did not expose tissue and then cooled for 15 minutes. The slides were then rinsed in deionised water and Tris-buffered saline, 0.05% Tween® 20 Detergent (TBST) wash solution. The tissue on the slides was encircled with a hydrophobic barrier pen and 200μL protein blocking solution (Akoya) was applied for 10 minutes and then replaced with the primary antibody. The desired antibody concentration was diluted in the blocking solution and the slides were incubated in a moist chamber at room temperature for one hour. The slides were rinsed in TBST and then 3 washes of 2 minutes each in TBST, this removed the blocking solution and any unbound antibody on the tissue. Followed by 200μL Opal Polymer Secondary antibody (Horseradish Peroxidase Mouse and Rabbit antibody), which was applied for 10 minutes onto the slides. This secondary antibody facilitates increasing the sensitivity and signal amplification of the 200μL Opal fluorophore, which was diluted in an amplification diluent (Akoya Biosciences). The Opal fluorophore was applied for 10 minutes and then removed with repeated TBST washes. Antigen stripping was then performed in the same method as antigen retrieval. These steps were repeated for every antibody, however, the final round of staining AR6 was used before the application of spectral DAPI or Opal 780. Following microwave treatment, 200μL of Opal 780 was pipetted onto each slide and incubated at room temperature for one hour. This was then washed off and DAPI was applied for 5 minutes at room temperature and then washed with TBST and deionised water. The slides were mounted with ProLong® diamond antifade mountant (Thermofisher) and coverslips and were then stored at room temperature before analysis. The approach to multiplex imaging involved cycles of antibody incubation with tissue, imaging and fluorophore deactivation as described previously. Whole slide scans of the stained mIHC slides were scanned using the PhenoImager™ HT (Akoya Biosciences) at 20 × magnification and saved as a QPTIFF file. Regions of interest (ROI) marked by IB were identified and stamped using x20 objective then imported into the InForm Analysis software (Akoya Biosciences) for unmixing and assessment of the stain quality using an algorithm that was created in the inForm® Tissue Analysis software (2.6 version, Akoya Biosciences). Each ROI was spectrally unmixed using a project-specific spectral library created using single-channel dyes and an autofluorescence control

### Dataset 3

Cancer metastasis in liver - Antibody panel is attached as Supplementary Table 3.

Multiplexed images were acquired using the Cell DIVE platform, as previously described.

### Dataset 4

Rheumatoid arthritis and osteoarthritis - Antibody panel is attached as Supplementary Table 4.

Multiplexed images were acquired using the Cell DIVE platform, as previously described.

### Dataset 5

PDAC - Antibody panel is attached as Supplementary Table 5.

Multiplexed images were acquired using the Cell DIVE platform, as previously described.

### Quality control

*Segmentation error and artifacts removal*

Cells with nuclear-to-cell ratio larger than 1 would be labelled as abnormal and will be removed. For markers in each cycle, if the exclusive markers are found expressed in the same region with extremely bright signals, users could set thresholds to label these cells by double positivity. If the markers with artifacts area in the same cycle are not exclusive, thresholds will be set by abnormal bright regions. Customed function based on DBSCAN will be used to connect and filter out artifact-affected region. More details on parameters selection will be found at the Jupyter notebook templates.

### Cell annotation

#### 1. Automated cell annotation

For each marker, we aimed to identify and normalize the data relative to a ’negative’ cell population, characterized by low or negligible marker expression. We employed a two-component GMM (GaussianMixture from scikit-learn, n_components=2) to model the distribution of marker intensities as a mixture of two Gaussian distributions, representing putative ’positive’ and ’negative’ cell populations. The GMM estimates the parameters of these two components: the means (μ₁ and μ₂), standard deviations (σ₁ and σ₂), and weights (w₁ and w₂). The weights represent the proportion of cells assigned to each component.

To accurately identify the ’negative’ cell population, we implemented a robust approach. First, we compared the weights of the two components. If one weight (w_max) was significantly larger (at least four times greater) than the other (w_min), we assigned the component with the larger weight as the ’negative’ population. This approach prioritizes the identification of a potentially large but low-expressing population. Following the identification of the ’negative’ population (with mean μ_neg and standard deviation σ_neg), we normalized the marker intensities (x) using a Z-score transformation: x(normalized) = (x−μneg )/σneg.

This transformation centres the ’negative’ population around 0 and scales the data so that the ’negative’ population has a standard deviation of 1, facilitating consistent comparison across markers and images. After normalization, we used the SCIMAP package (v2.0.0) in conjunction with the napari image viewer to define marker-specific thresholds/gates for cell phenotyping. Specifically, we visualized the normalized marker intensities using napari and interactively defined thresholds that effectively separated positive and negative cell populations based on the normalized distributions. These thresholds were then applied across all cells in the dataset using SCIMAP functions.

### 2. SVM based cell annotation

A Support Vector Machine (SVM) classifier (using scikit-learn v1.6.0 with a RBF kernel was trained to predict cell phenotypes. Half of the dataset was randomly selected as the training set, with the initial SCIMAP-derived cell annotations serving as the target labels. The model’s performance was evaluated on the remaining half of the data using accuracy as the primary metric. The trained SVM model was then applied to the entire dataset to generate probability scores for each cell belonging to each of the defined phenotypes. These probability scores were visualized using UMAP embedding with parameters n_neighbours=10, min_dist=1. Potential annotation errors were identified by defining outlier cells based on a probability threshold for their assigned phenotype (e.g., tumour cells with a PCK+ probability < 0.9). Cells in transitional states were investigated by examining the probability distributions across multiple related phenotypes within defined cell populations.

### Density distribution

#### General Density

To quantify the spatial distribution of cell populations across whole-slide images, we implemented a tile-based density analysis. Each image was systematically overlaid with a grid of non-overlapping square tiles with a side length of 128 pixels. This tile size was chosen to capture local variations in cell density while maintaining computational feasibility across large images and can be adjusted by the user depending on the biological context and desired resolution. For each tile within an image, we calculated two distinct measures of general cell density using a custom Python function (calculate_density). This function takes a DataFrame (fov_table) containing single-cell information within the tile and the total number of pixels in the tile (128×128=16384 pixels) as input. Pixel-based Density: To account for variations in cell size, we calculated the total area (in pixels) occupied by all segmented cells within the tile. This total pixel area was then divided by the total number of pixels in the tile and multiplied by 100 to represent the percentage of the tile area occupied by cellular material. Object-based Density: To quantify the overall cellularity of each tile, we determined the total number of unique segmented cells present within the tile. This count was then divided by the total number of pixels in the tile and multiplied by 100 to represent the percentage of cells per total pixels in the tile. These density values were subsequently used for visualization by overlaying them onto the original images, as described in the Results section.

### Local density

To assess the spatial distribution of cells at a finer scale, we calculated the local density for each individual cell. For each image and each identified cell phenotype within that image, we utilized a k-nearest neighbours (kNN) approach. Specifically, for each cell of a given phenotype, we employed the BallTree algorithm (from scikit-learn) to identify its k nearest neighbors based on their centroid coordinates (’X_centroid’, ’Y_centroid’). The number of neighbors, k, was set to 5 (or k=min (6, number of cells of that phenotype) to avoid issues with small populations). The local density for each target cell was then defined as the inverse of the mean Euclidean distance to its k nearest neighbours:

This calculation was performed separately for each cell within each phenotype and each image. The resulting local density values were stored as a new feature (’density’) in the object’s observation metadata (adata.obs), allowing for the analysis and visualization of local clustering patterns across different cell populations and tissue regions. The mean distance to the k nearest neighbours (’mean_dist’) was also stored for potential further analysis.

### Boundary identification

Cells are categorized into “cell-of-interest” and “other cells”, which would be the input of the boundary identification module. “cell-of-interest” located closer with each will be clustered together using customed function based on DBSCAN. ConcaveHull will be used to identify the boundary cells. More details on parameters selection will be found at the Jupyter notebook templates.

### Pseudotime analysis

*Epithelial cell clusters identification.* The “cell-of-interest” here is defined as epithelial cells (CK19+ cells). Based on the distribution of the epithelial cells, closely located epithelial cells would be clustered together and assigned a unique cluster number using the customed function based on DBSCAN.

*Generating cluster morphology information*.

1. *Computing Proliferating Index.* Proliferating index = number of Ki67+ epithelial cells / total number of the epithelial cluster.
2. *Computing the cluster level morphological features*. To quantify morphological heterogeneity among epithelial cell clusters, we developed an automated analysis pipeline to extract and calculate shape-based features from binary cell segmentation masks for each cluster based on segmentation label maps. The resulting binary mask was refined by removing small holes (area threshold of 20 pixels) and applying binary morphological closing, which were then used to compute the medial axis ^1^, a skeletonized representation of the shape, along with the corresponding distance transform, using the “medial_axis” function from the skimage.morphology module. The distance values along the medial axis were doubled to estimate local thickness, and all measurements were rescaled to account for the image downsampling factor (0.5×). The following *thickness-based morphological metrics* were calculated:
  a. *Maximum thickness*: the largest local thickness value within the cluster.
  b. *Mean thickness:* the average of all local thickness values along the medial axis.
  c. *Median thickness:* the median of local thickness values, providing a robust central tendency measure.
  d. *Standard deviation of thickness:* used to assess intra-cluster morphological variability
  Other cluster level morphological properties were computed using regionprops ^1^ to the refined cluster segmentation masks. For each epithelial cell cluster, we extracted the following metrics:

e. *Centroid coordinates (x, y):* spatial centre of the cluster.
f. *Area and perimeter:* size and boundary length of the region.
g. *Eccentricity:* elongation of the region based on fitted ellipses.
h. *Convexity:* ratio of convex hull perimeter (computed by scipy.spatial.ConvexHull) to actual perimeter.
i. *Compactness and circularity:* shape regularity, calculated as 4π*area/perimeter^2^.
j. *Elongation:* ratio of major to minor axis lengths.
k. *Fractal dimension:* complexity of the boundary, estimated as log10(boundary pixels)/log10(area)
l. *Orientation:* angle of the major axis of the region.
m. *Radial spatial features:* including the average radial distance of all cells from the cluster centroid, variance in cell layer density (10 radial bins), variance in radial thickness, and spatial entropy of cell distribution.
3. *Computing the cell level morphological features.* To quantify cell morphology, we extracted a set of morphological features at the single-cell level. These features were either computed directly using Pixie or derived from additional image-based calculations. The resulting features were subsequently grouped by epithelial clusters, and summary statistics were computed for each cluster to support downstream pseudotime analysis.

a. *Features extracted from Pixie* The following features were obtained directly from Pixie: *area, eccentricity, major_axis_length, minor_axis_length, perimeter, convex_area, equivalent_diameter, major_minor_axis_ratio, perim_square_over_area, major_axis_equiv_diam_ratio, convex_hull_resid, centroid_dif, num_concavities, nc_ratio*
b. *Additional features* Other relevant morphological features were computed using skimage.measure.regionprops and custom calculations:

i. *orientation, solidity, feret_diameter_max:* Extracted using regionprops.
ii. *Circularity:* a shape descriptor quantifying how close the cell shape is to a perfect circle, calculated as: (4π*area)/perimeter^2^.
iii. *Fractal_dimension:* an approximation of the complexity of the cell boundary, defined as: log(perimeter)/log(area).
iv. *Boundary_irregularity:* a measure of how irregular the cell boundary is relative to its convex hull: perimeter/√convex area
4. *Computing the pixel level morphological features*. To better capture the transition of normal epithelial cells to malignant cells, we quantified a set of pixel-level morphological features to assess changes in cell polarity and nuclear texture. These features were subsequently incorporated into our spatialtemporal pseudotime analysis. Each feature was computed per cell using a parallelized image-processing pipeline in Python. For computational efficiency, each cell’s local image region was cropped with a 2-pixel padding around its segmentation mask. The following features were extracted:

a. *Polarity score:* polarity was quantified as the normalized Euclidean distance between the geometric centroid of the cell mask and the intensity-weighted centroid of a membrane marker channel (e.g., CK19). A low polarity score indicated symmetric distribution, while a higher score reflects asymmetric localization. The polarity score is calculated as: *||geo_centroid – intensity_centroid|| / area*.
b. *Moment of Inertia:* the spatial moment of inertia around the intensity centroid was calculated to reflect how widely membrane marker signal is dispersed. Higer values indicate more diffuse signal distributions, suggesting disrupted polarity or membrane disorganization. The moment of inertia is calculated as: *∑(distance² × intensity) from intensity centroid*
c. *Intensity ratio:* to assess the relative abundance of two membrane markers, we computed the ratio of mean intensities between 2 channels (e.g., NaKATPase / CK19) per cell.
d. *Pearson correlation Coefficient (PCC):* PCC was calculated between pixel intensities of 2 membrane markers within the cell mask to evaluate the spatial co-localization of these signals. A high PCC indicates a high degree of overlap, while a low PCC suggests spatial separation or differential expression.
e. *Shannon entropy (nuclear texture):* Shannon entropy was calculated from the DAPI channel to capture intensity randomness, reflecting chromatin condensation status. Higher entropy values indicate a more disorganized, decondensed nuclear texture, usually associated with malignancy or transcriptional activation.
f. *Lacunarity (chromatin texture heterogeneity):* we used a sliding-window-based approach to quantify lacunarity, a measure of texture heterogeneity in the DAPI channel. Larger lacunarity values suggest increased chromatin irregularity or fragmentation.
g. *Haralick texture features:* 4 haralick features were extracted from the DAPI channel using the Gray-Level-Co-occurrence Matrix (GLCM): contrast, correlation, energy, and homogeneity. These features capture spatial relationships of pixel intensities, revealing fine-scale nuclear texture variations associated with neoplasia.

#### Measuring the surrounding niche cell type composition

To assess the cellular context surrounding each epithelial cluster, we first identified the boundary for each epithelial cluster using customed function. A peri-epithelial niche was then defined as the region extending 30 μm outward from the cluster boundary. This expansion was applied isotropically. All single cells whose centroids fell within this 30 μm boundary expansion were considered part of the niche. The cell type composition within each niche was subsequently quantified based on prior single-cell phenotyping, allowing for detailed characterization of the immune and stromal landscape associated with each epithelial cluster.

#### Data transformation and pseudotime trajectory inference

For each epithelial cluster, morphological parameters and the corresponding peri-cluster cell type composition were combined into a unified feature matrix. Prior to analysis, the distribution of each feature was evaluated, and appropriate transformations were applied. All features were then standardized to ensure comparability across scales.

To visualize the global structure of the data and facilitate trajectory inference, dimensionality reduction was performed using Uniform Manifold Approximation and Projection (UMAP). Pseudotime progression of epithelial clusters was subsequently inferred with ElPiGraph.

### Weighted Feature Correlated Network Analysis (WFCNA)

A feature matrix was generated with 877 spatial metrics across cell types, 22 metrics across neighbourhoods, 56 metrics across ECM. Detailed list of features with their cluster assignment is provided in supplementary materials **(Figure_7_Suppl)**. Features with >40% missing data or <8 unique values were filtered. Remaining missing data were imputed using the missForest R package with 20 iterations. Features with significantly altered means post-imputation were excluded. A subsample depth was calculated based on minimum feature coverage. A correlation matrix was then generated using a subsampling approach. For each pair of features, a random subsample of minimum sample observations was drawn from the available data points for both features. The Pearson correlation coefficient was calculated for this subsample. This process was repeated 10,000 times for each feature pair, and the mean of these 10,000 correlation coefficients was assigned as the final correlation value between the features. Features were clustered into modules using the dynamicTreeCut R package enabling the identification of robust and biologically meaningful modules. Type-specific parameters for top_clust, bot_clust, and min_size were also applied: for cell type data, top_clust = 45, bot_clust = 25, and min_size = 15; for ECM data, top_clust = 10, bot_clust = 5, and min_size = 3; and for neighbourhood data, top_clust = 7, bot_clust = 3, and min_size = 2. After features were clustered into modules, a representative value for each module, first principal component (PC1), was calculated. This was achieved using the prcomp function on the scaled and imputed feature matrix (scaled_matrix). For each module, the first principal component was extracted, which optimally explains the variance within that module and used for downstream analysis of RA and OA images.

## Code and data availability

All code is available via https://github.com/Bashford-Rogers-lab/SpatioEv. Raw imaging data will be available upon reasonable request.

## Acknowledgements

S.W. is supported by Cancer Research UK. S.A. is funded by St John’s Clarendon Fund scholarship. E.C is supported by NIHR BRC-Oxford. J.B.R. was supported by Versus Arthritis and C.M. by the MRC.

We acknowledge the contribution to this study made by the Oxford Centre for Histopathology Research and the Oxford Radcliffe Biobank, which are funded by the University of Oxford, the Oxford CRUK Cancer centre, the NIHR Oxford Biomedical Research Centre (BRC) (Molecular Diagnostics Theme/Multimodal Pathology Subtheme and the NIHR CRN Thames Valley network.

## Authors’ contributions

S.W., S.A., K.M. and R.B.-R. conceived and designed the analysis. S.W., C.L., J.B.R, N.M, C.M., E.C. collected the data and/or samples. S.W., R.B.-R, S.A, L.O. contributed to code development. S.W., S.A. and R.B.-R. performed the analyses. All authors contributed intellectual input/interpretation. S.W., S.A., N.M., and R.B.R. wrote the paper with input from all other authors. S.W. and S.A. contributed equally. K.M. and R.B.-R. jointly supervised this work.

## Ethics approval and consent to participate

Informed consent was obtained for all patients. The study was in strict compliance with all institutional ethical regulations. PDAC Surrey Cohort: PDAC Formalin-fixed paraffin embedded (FFPE) tissue was retrieved with ethical approval (study IRAS project 277406, ethical reference: 20/SW/0105).

HNSCC tissue was retrieved with ethical approval (study IRAS project 262470, ethical reference: 19/SC/0173)

Need to add for RA vs OA.

## Supplemental Figures

**Supplementary Figure 1.**
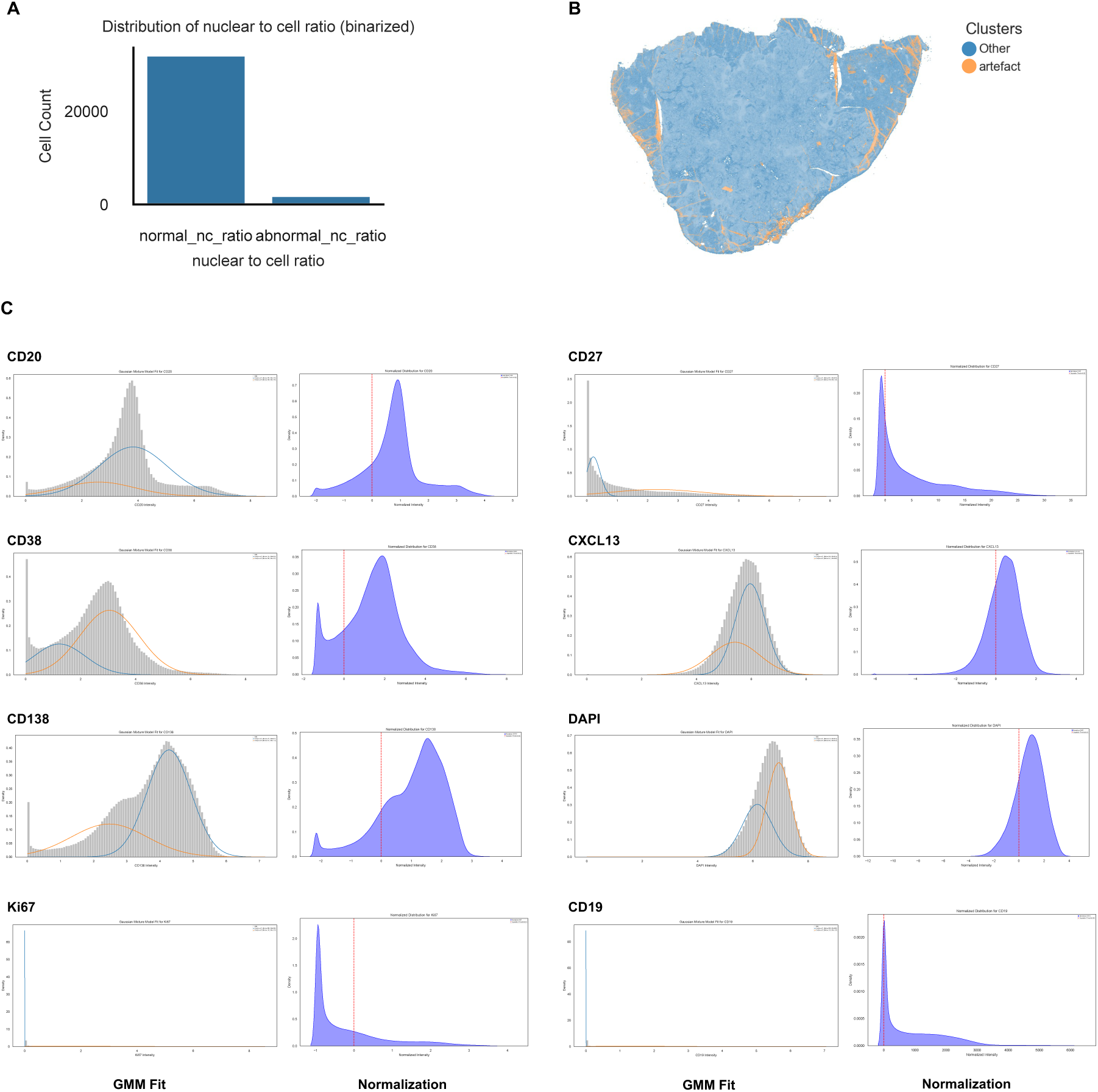
Quality control. **A.** Bar plot showing the binarized distribution of normal and abnormal Nuclear-to-cell ratio of the cell segmented on the pancreatic ductal adenocarcinoma (PDAC) image. **B.** Scatter plot of colorectal cancer liver metastasis image colored by “Other cells” and “artefact cells”. **C.** (Left) Distribution of marker intensity across cells and the fitted Gaussian Mixture Model (GMM). The grey bars represent the histogram of the raw intensity data. The solid blue and orange lines represent the two Gaussian components identified by the GMM. Each component corresponds to a distinct subpopulation of cells. (Right) Distribution of marker intensity after normalization using the Gaussian Mixture model.

**Supplementary Figure 2.**
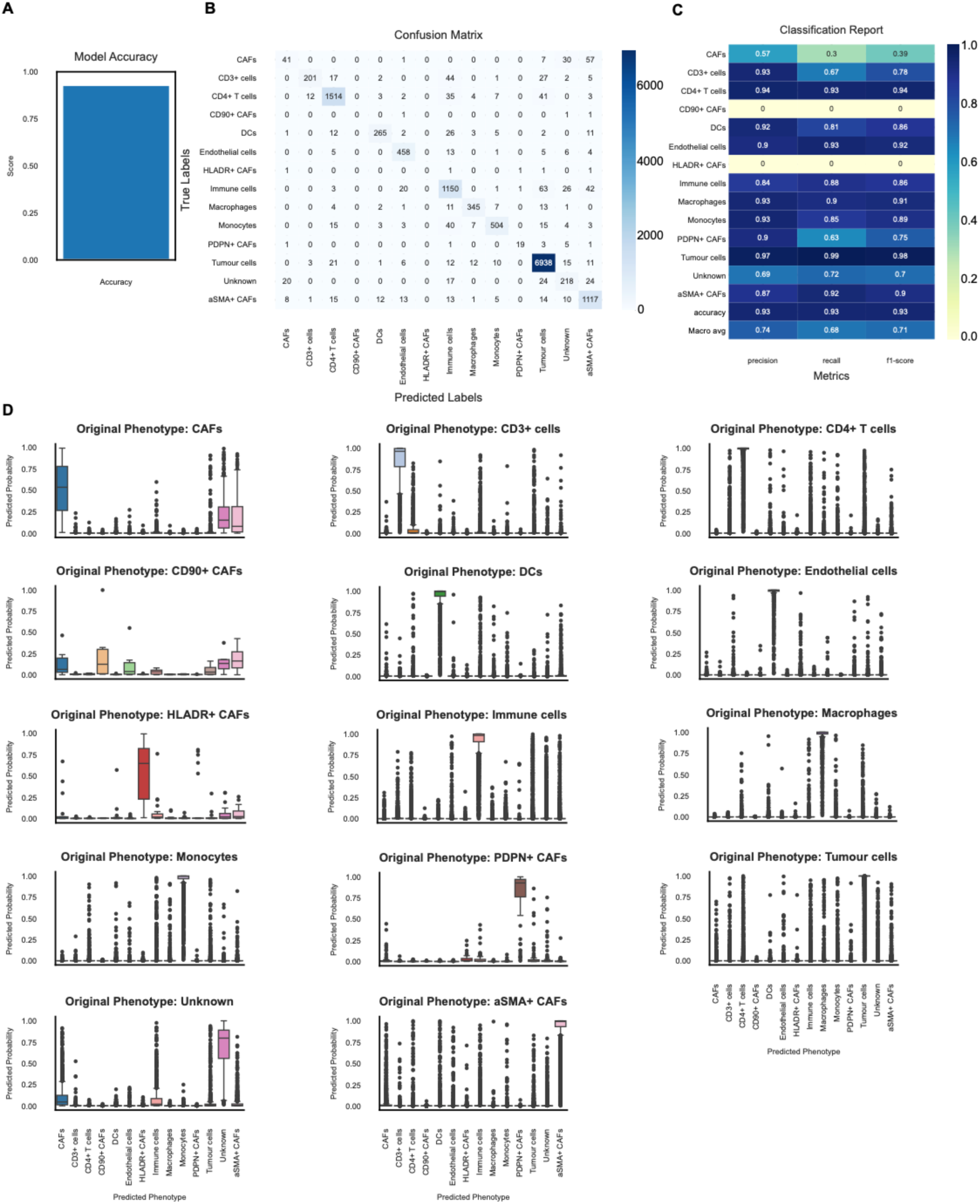
SVM-based approach for annotation improvement. **A.** Bar plot showing the accuracy of SVM model predicting the phenotypes. **B.** Heatmap showing the confusion matrix of the SVM model. **C.** Heatmap showing the classification report of the SVM model. **D.** Boxplots showing the distribution of predicted probabilities of belonging to each phenotype for originally annotated populations.

**Supplemental Figure 3.**
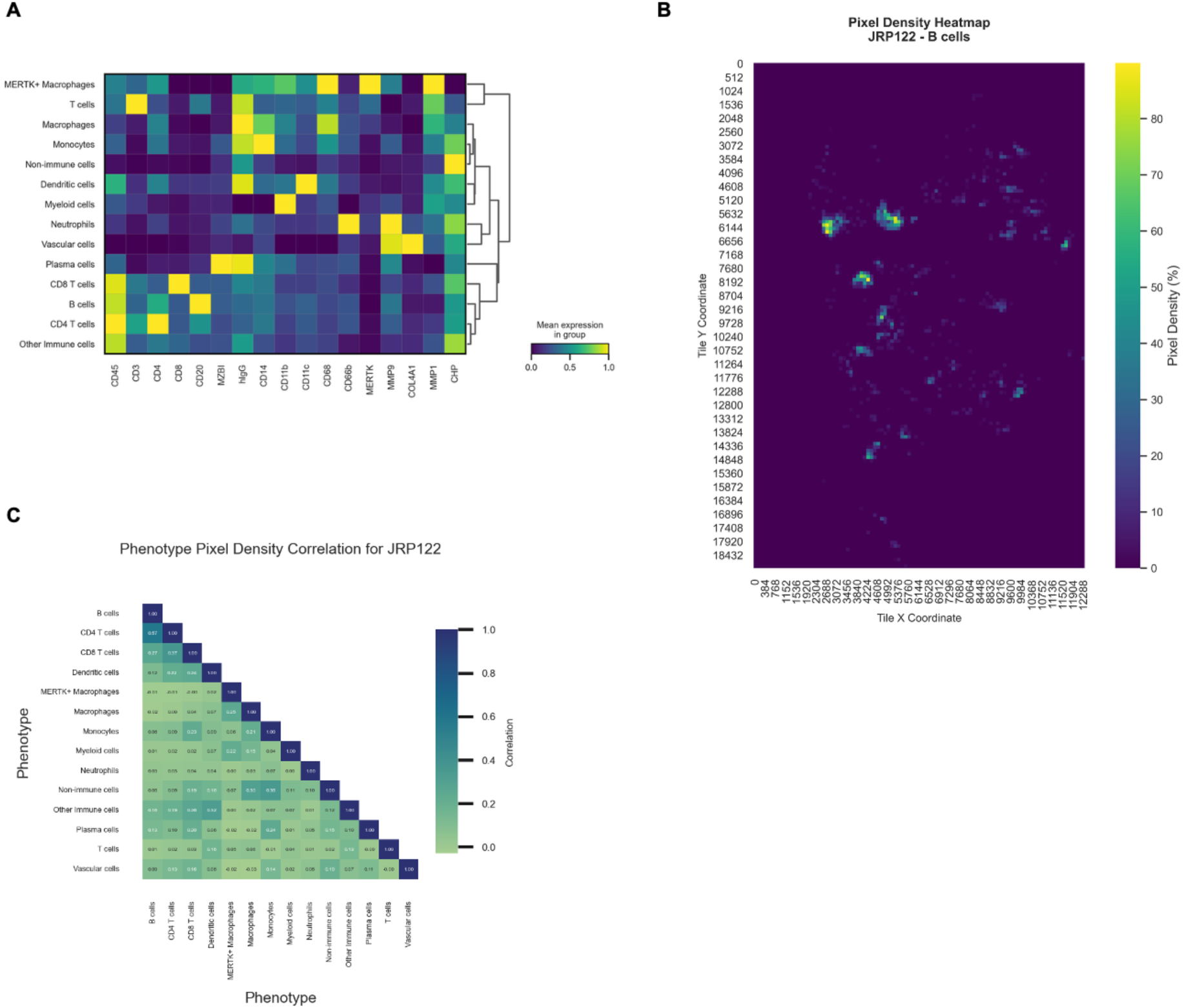
**A.** Heatmap showing the annotation of RA and OA image cohort. **B.** Representative RA image colored by the pixel density of B cells. **C**. Pearson correlation matrix of pixel density across different cell phenotypes.

**Supplementary Figure 4.**
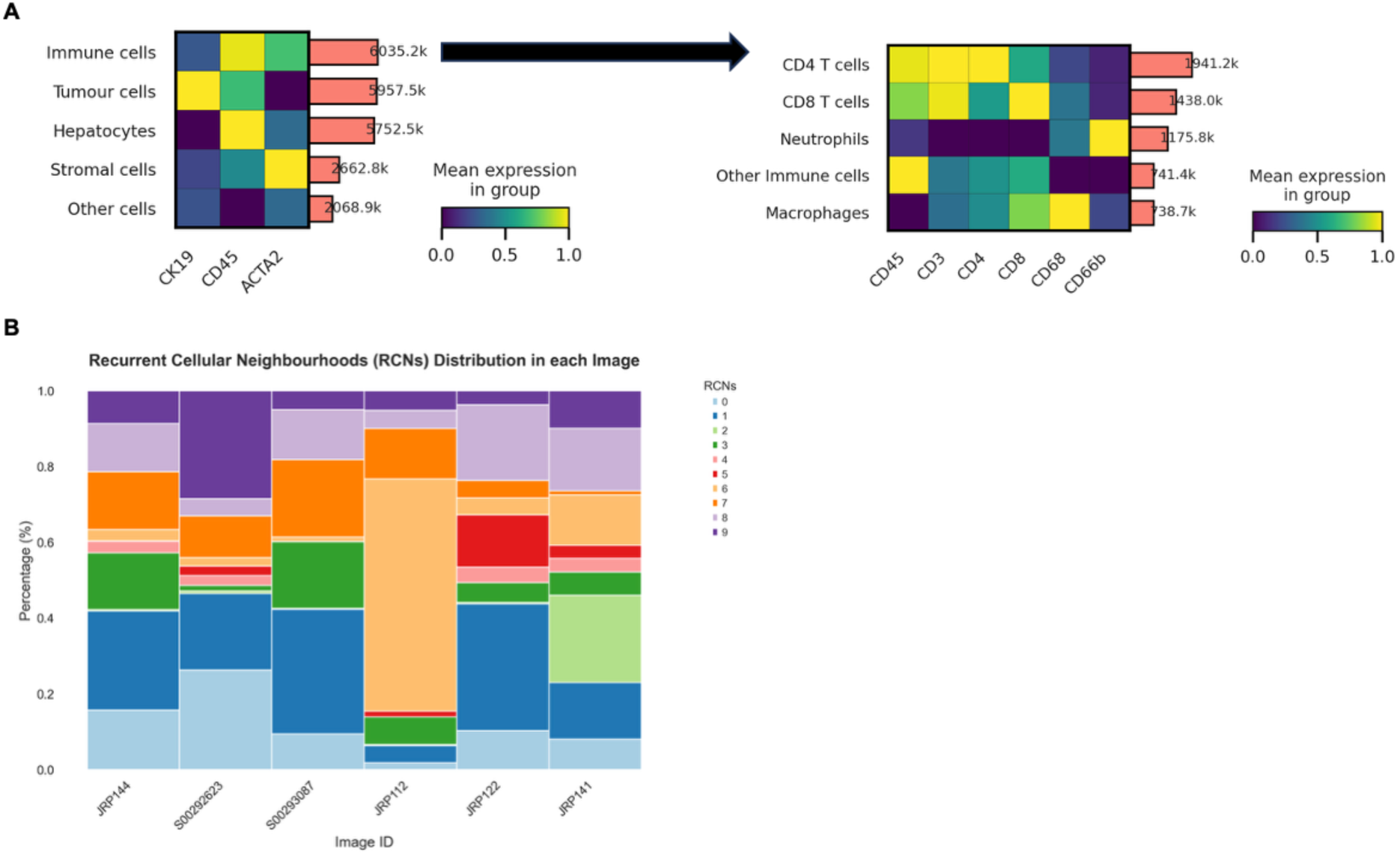
Automated ROI selections RA and OA image cohort. **A.** Heatmap showing the annotation of the RA and OA images at the general cell type level and immune cell type level. **B.** The distribution of RCNs across RA and OA images.

**Supplementary Figure 5.**
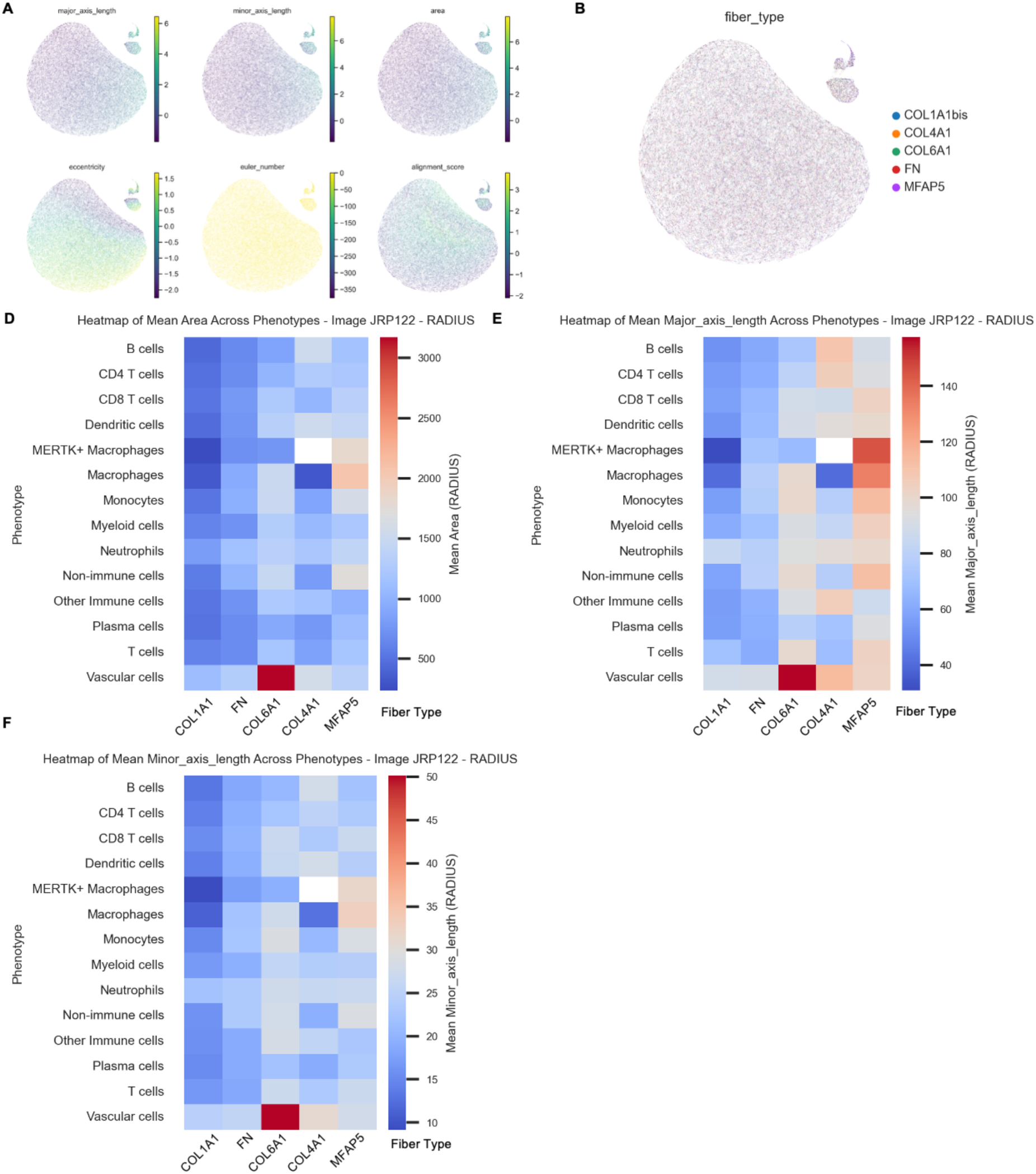
Integrated spatial analysis of extracellular matrix fibers and cellular features. **A.** UMAP plots of extracellular matrix fibers colored by “major axis length”, “minor axis length”, “area”, “eccentricity”, “Euler number”, and “alignment score”. **B.** UMAP plot of extracellular matrix fibers colored by fiber types. **C.** Heatmap of mean area of fibers surrounding each cell types. **D.** Heatmap of mean major axis length of fibers surrounding each cell types. **E.** Heatmap of mean minor axis length of fibers surrounding each cell types.

**Supplementary Figure 6.**
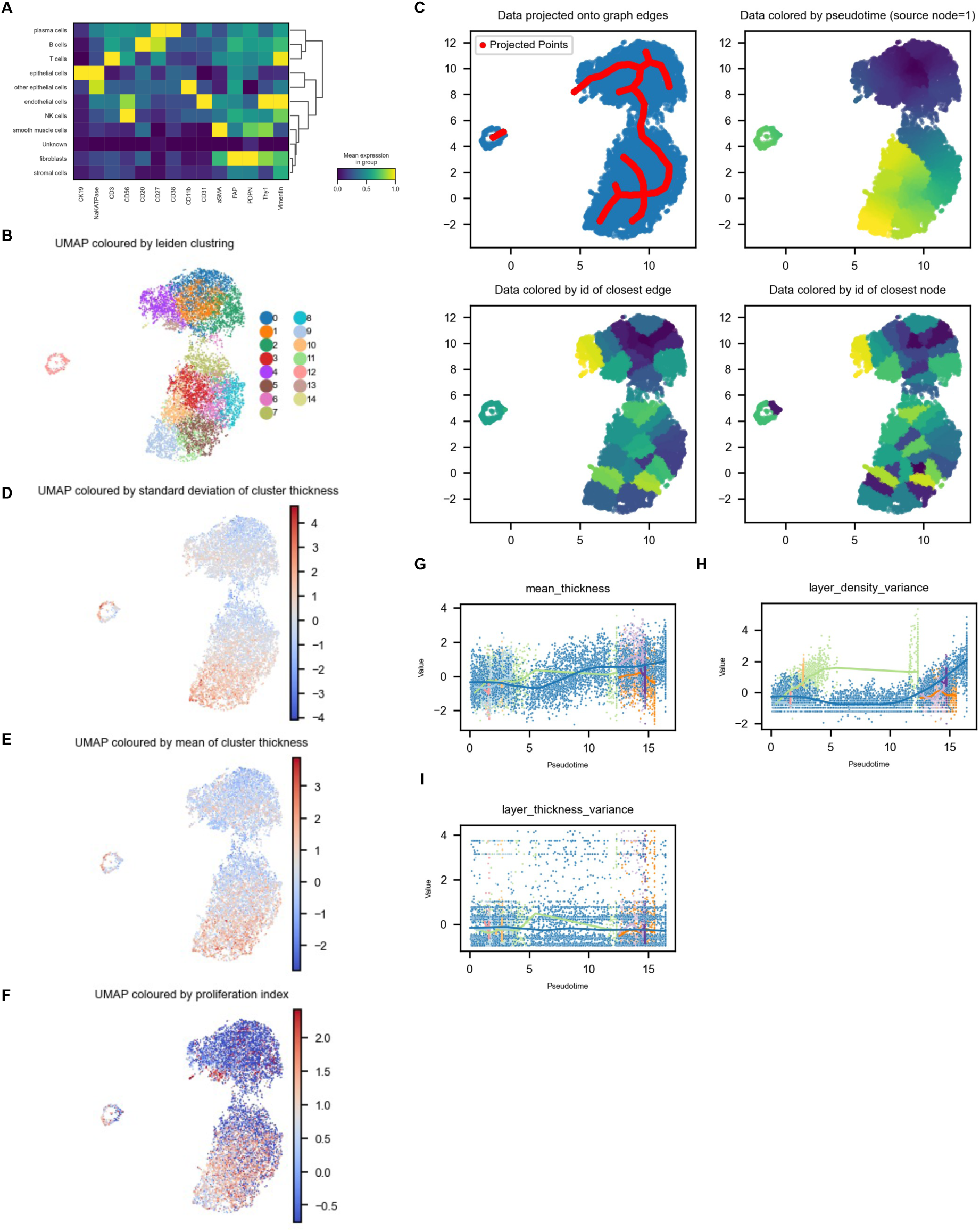
Spatiotemporal Trajectory Inference Reveals Divergent Evolutionary Pathways and Dynamic Niche Composition in Pancreatic Ductal Adenocarcinoma. **A.** Heatmap illustrating protein signatures for identified cell types on the PDAC image. **B.** UMAP of epithelial clusters colored by Leiden clustering. **C.** UMAP of epithelial clusters overlayed with ElPiGraph projected points (upper left); UMAP of epithelial clusters colored by pseudotime values (upper right); UMAP of epithelial clusters colored by id of closest edge (bottom left); UMAP of epithelial clusters colored by id of closest node. **D-F.** UMAP of epithelial clusters colored by standard deviation of cluster thickness (D), mean of cluster thickness (E), and proliferation index (F). **G-I.** Trends of cluster-level parameters: mean thickness (G), layer density variance (H), layer thickness variance (I), plotted over pseudotime, colored by branch ids.

**Supplementary Figure 7.**
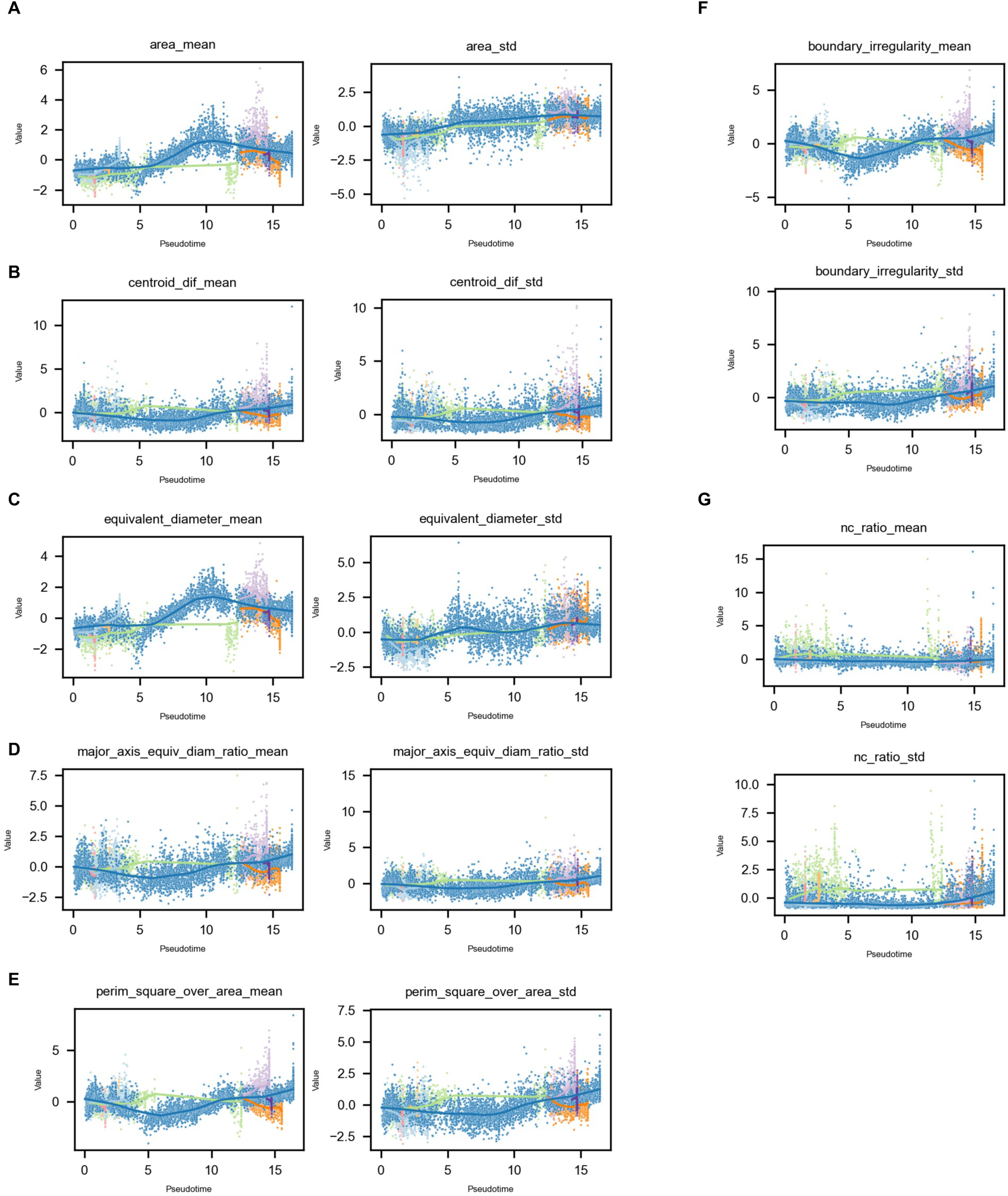
Spatiotemporal Trajectory Inference Reveals Divergent Evolutionary Pathways and Dynamic Niche Composition in Pancreatic Ductal Adenocarcinoma. **A-G.** Trends of cell-level parameters: mean and standard deviation of area (A), mean and standard deviation of centroid difference (B), mean and standard deviation of equivalent diameter (C), mean and standard deviation of major axis equivalent diameter ratio (D), mean and standard deviation of perimeter square over area (E), mean and standard deviation of boundary irregularity (F), mean and standard deviation of nuclear-to-cell ratio (G), plotted over pseudotime, colored by branch ids.

**Supplementary Figure 8.**
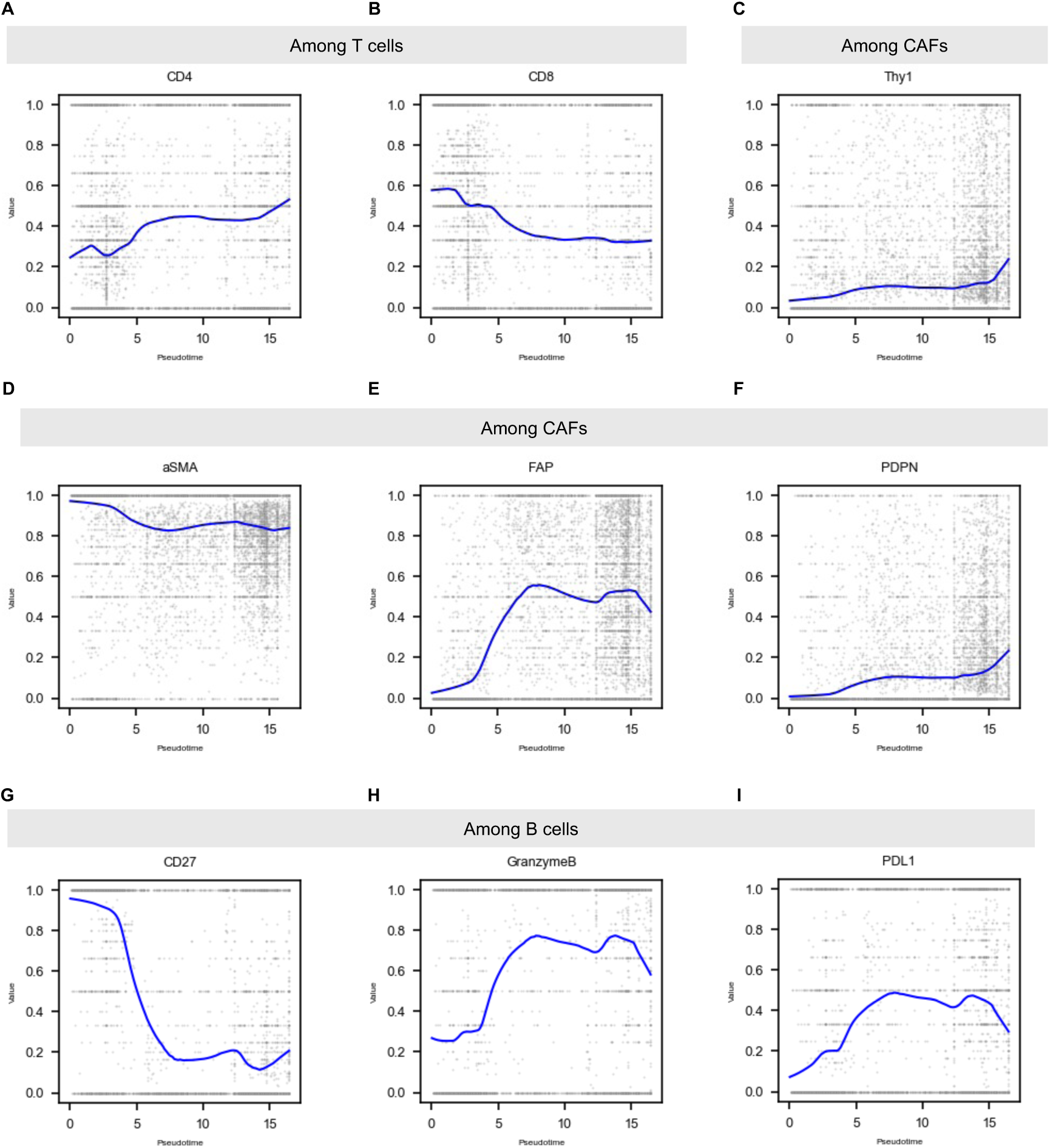
Spatiotemporal Trajectory Inference Reveals Divergent Evolutionary Pathways and Dynamic Niche Composition in Pancreatic Ductal Adenocarcinoma. **A-B.** Trends of CD4 (A) and CD8 (B) positivity among T cells (CD3+) over pseudotime. **C-F.** Trends of Thy1 (C), aSMA (D), FAP (E), PDPN (F) positivity among CAFs over pseudotime. **G-I.** Trends of CD27 (G), GZMB (H), PDL1 (I) positivity among B cells over pseudotime.

**Supplementary Figure 9.**
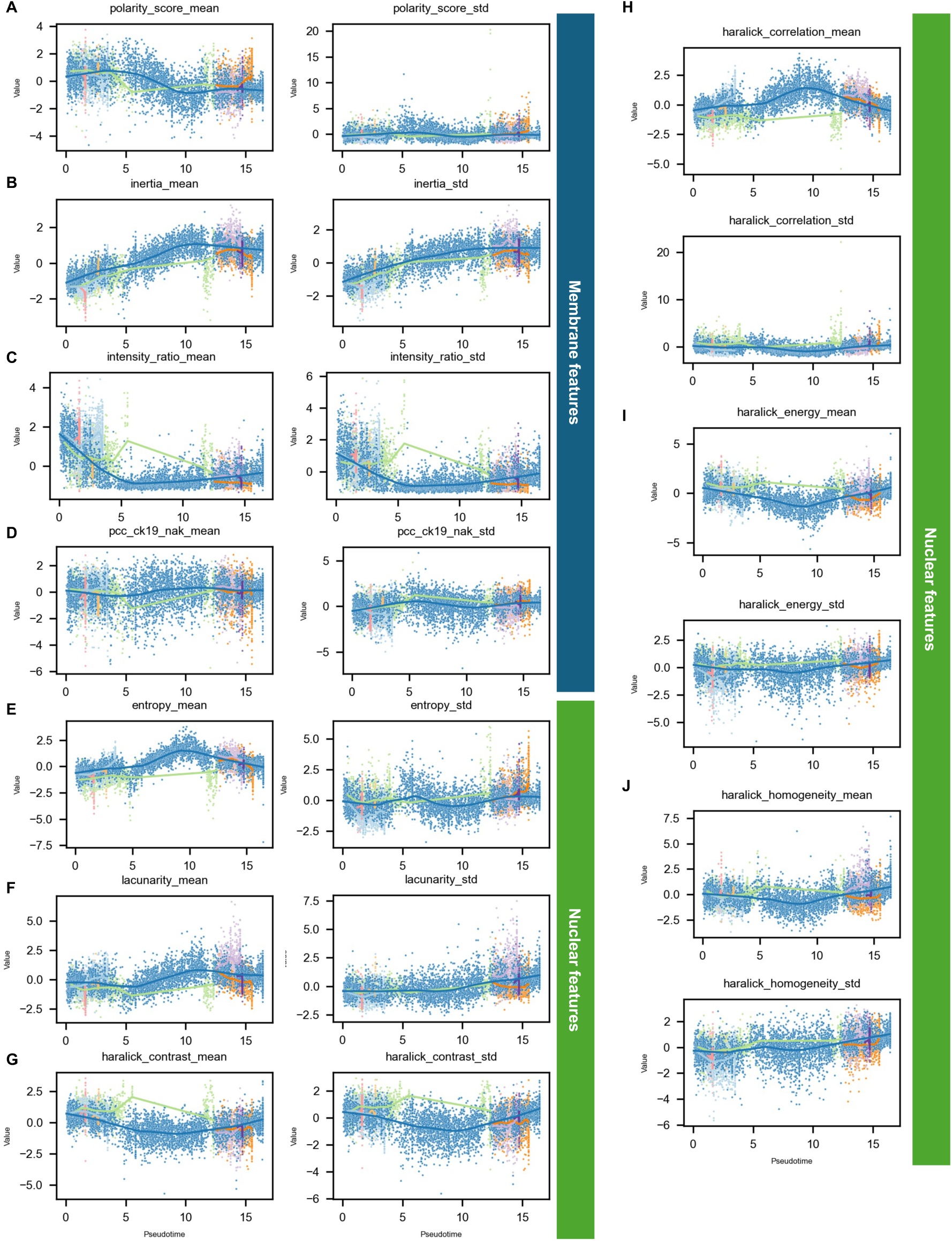
Spatiotemporal Trajectory Inference Reveals Divergent Evolutionary Pathways and Dynamic Niche Composition in Pancreatic Ductal Adenocarcinoma. **A-D.** Trends of pixel-level membrane features: polarity score (A), inertia (B), intensity ratio (C), Pearson correlation coefficient of CK19 and NaKATPase (D), over pseudotime, colored by branch ids. **E-J.** Trends of pixel-level nuclear features: entropy (E), lacunarity (F), haralick contrast (G), haralick correlation (H), haralick energy (I), haralick homogeneity (J), over pseudotime, colored by branch id.

**Supplementary Table 1.**

| **Antibody** | **Catalog** | **Clone** | **Company** | **Concentration** |
| --- | --- | --- | --- | --- |
| IGFBP5 |  |  |  |  |
| COL4A1 | 51-9871-82 | 1042 | Invitrogen | 5 µg/ml |
| CD146 | ab196448 | EPR3208 | Abcam | 5 µg/ml |
| RS6 | sc-74459 | C-8 | SCBT | 5 µg/ml |
| CD31 | NBP2-33154 | C31.3 | Novus | 5 µg/ml |
| Vimentin | ab185030 | EPR3776 | Abcam | 5 µg/ml |
| CD68 | ab280860 | EPR20545 | Abcam | 5 µg/ml |
| CD19 | ab196515 | EPR5906 | Abcam | 5 µg/ml |
| PCK | ab6410 | PCK-26 | Abcam | 5 µg/ml |
| CD3 | ab245731 | SP162 | Abcam | 5 µg/ml |
| NaKATPase | ab198367 | EP1845Y | Abcam | 5 µg/ml |
| COL1A1 | 72026 | E8F4L | CST | 5 µg/ml |
| CD90 | ab307267 | EPR3132 | Abcam | 5 µg/ml |
| PDPN | 916610 | D2-40 | BioLegend | 5 µg/ml |
| CK19 | ab205445 | EPR1579Y | Abcam | 5 µg/ml |
| Tenascin C | ab271877 | EPR4219 | Abcam | 5 µg/ml |
| COL6A1 | ab182744 | EPR17072 | Abcam | 5 µg/ml |
| aSMA | ab184675 | 1A4 | Abcam | 5 µg/ml |
| Fibronectin | ab275110 | F1 | Abcam | 5 µg/ml |
| CD45 | ab282747 | RM1007 | Abcam | 5 µg/ml |
| HLADR | sc-53319 | TAL 1B5 | SCBT | 5 µg/ml |
| CD11c | ab279329 | EP1347Y | Abcam | 5 µg/ml |
| CD4 | ab196147 | EPR6855 | Abcam | 5 µg/ml |
| CD14 | ab226121 | EPR3653 | Abcam | 5 µg/ml |

**Supplementary Table 2.**

| Antigen | Antibody | Cell/Marker of Interest | Position | Primary antibody concentration | Opal fluorophore | Opal concentration |
| --- | --- | --- | --- | --- | --- | --- |
| CXCL13 | PA5-28827 | Promotes the migration of B cells | 1 | 1:150 | 480 | 1:100 |
| CD138 | DL-101 | Precursors of B cells | 2 | 1:500 | 650 | 1:50 |
| Ki67 | MA5-14520 | Neuronal marker of cell cycling and proliferation. | 3 | 1:2000 | 540 | 1:150 |
| CD38 | 25-0389-42 | Plasma cells and activated T lymphocytes | 4 | 1:300 | 690 | 1:100 |
| CD19 | ZR212 | B cell proliferation | 5 | 1:600 | 570 | 1:100 |
| CD27 | LG.7F9 | Memory B cells and mature T lymphocytes | 6 | 1:3000 | 620 | 1:100 |
| CD20 | PA5-16701 | Mature B cell proliferation | 7 | 1:600 | 520 | 1:150 |
| Pan-CK | AE1/AE3 | Epithelial tissue marker | 8 | 1:300 | 780 | 1:25 |

**Supplementary Table 3.**
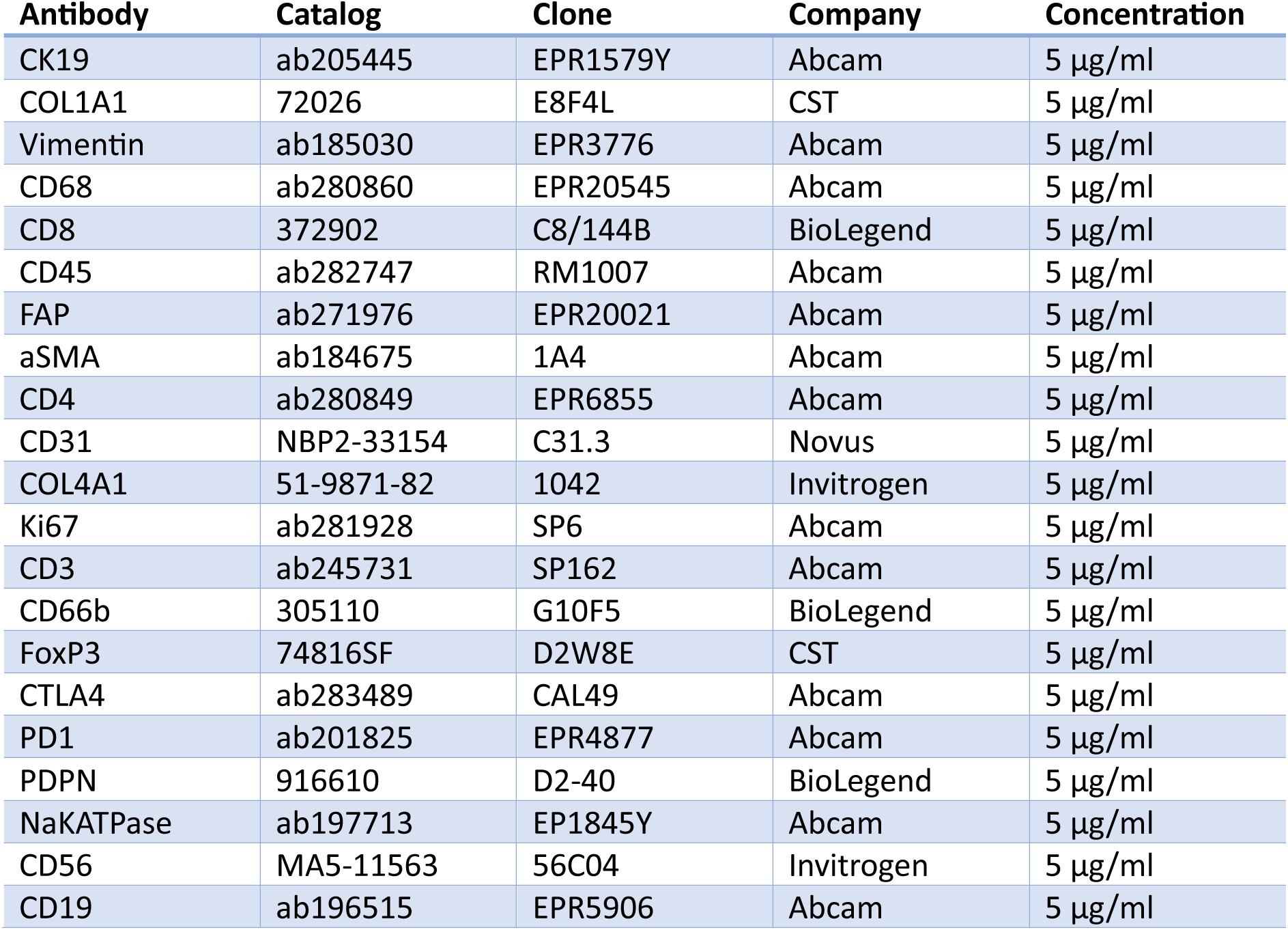

| **Antibody** | **Catalog** | **Clone** | **Company** | **Concentration** |
| --- | --- | --- | --- | --- |
| CK19 | ab205445 | EPR1579Y | Abcam | 5 µg/ml |
| COL1A1 | 72026 | E8F4L | CST | 5 µg/ml |
| Vimentin | ab185030 | EPR3776 | Abcam | 5 µg/ml |
| CD68 | ab280860 | EPR20545 | Abcam | 5 µg/ml |
| CD8 | 372902 | C8/144B | BioLegend | 5 µg/ml |
| CD45 | ab282747 | RM1007 | Abcam | 5 µg/ml |
| FAP | ab271976 | EPR20021 | Abcam | 5 µg/ml |
| aSMA | ab184675 | 1A4 | Abcam | 5 µg/ml |
| CD4 | ab280849 | EPR6855 | Abcam | 5 µg/ml |
| CD31 | NBP2-33154 | C31.3 | Novus | 5 µg/ml |
| COL4A1 | 51-9871-82 | 1042 | Invitrogen | 5 µg/ml |
| Ki67 | ab281928 | SP6 | Abcam | 5 µg/ml |
| CD3 | ab245731 | SP162 | Abcam | 5 µg/ml |
| CD66b | 305110 | G10F5 | BioLegend | 5 µg/ml |
| FoxP3 | 74816SF | D2W8E | CST | 5 µg/ml |
| CTLA4 | ab283489 | CAL49 | Abcam | 5 µg/ml |
| PD1 | ab201825 | EPR4877 | Abcam | 5 µg/ml |
| PDPN | 916610 | D2-40 | BioLegend | 5 µg/ml |
| NaKATPase | ab197713 | EP1845Y | Abcam | 5 µg/ml |
| CD56 | MA5-11563 | 56C04 | Invitrogen | 5 µg/ml |
| CD19 | ab196515 | EPR5906 | Abcam | 5 µg/ml |

**Supplementary Table 4.**

| **Antibody** | **Catalog** | **Clone** | **Company** | **Concentration** |
| --- | --- | --- | --- | --- |
| MMP2 |  |  |  |  |
| CHP |  |  |  |  |
| CD66b | 305110 | G10F5 |  | 5 µg/ml |
| COL3A1 | ab237238 | EPR17673 | Abcam | 5 µg/ml |
| MMP9 |  |  |  |  |
| CD14 | ab226121 | EPR3653 | Abcam | 5 µg/ml |
| COL1A1 | 72026 | E8F4L | CST | 5 µg/ml |
| COL12A1 |  |  |  |  |
| COL6A1 | ab182744 | EPR17072 | Abcam | 5 µg/ml |
| CD3 | ab307134 | SP162 | Abcam | 5 µg/ml |
| Fibronectin | ab275110 | F1 | Abcam | 5 µg/ml |
| CD20 | ab198943 | EP459Y | Abcam | 5 µg/ml |
| MERTK | ab307385 | Y323 | Abcam | 5 µg/ml |
| CD4 | ab280849 | EPR6855 | Abcam | 5 µg/ml |
| COL4A1 | 51-9871-82 | 1042 | Invitrogen | 5 µg/ml |
| CD45 | ab282747 | RM1007 | Abcam | 5 µg/ml |
| CD68 | ab280860 | EPR20545 | Abcam | 5 µg/ml |
| CD8 | 372902 | C8/144B | BioLegend | 5 µg/ml |
| IgG |  |  |  |  |
| CD11c | ab279329 | EP1347Y | Abcam | 5 µg/ml |
| MZBI |  |  |  |  |
| CD11b |  |  |  |  |
| LYVE1 |  |  |  |  |
| MFAP5 |  |  |  |  |
| MMP1 |  |  |  |  |

**Supplementary Table 5.**

| **Antibody** | **Catalog** | **Clone** | **Company** | **Concentration** |
| --- | --- | --- | --- | --- |
| CD8 | 372902 | C8/144B | BioLegend | 5 µg/ml |
| Ki67 | ab281928 | SP6 | Abcam | 5 µg/ml |
| CD31 | NBP2-33154 | C31.3 | Novus | 5 µg/ml |
| CD4 | ab280849 | EPR6855 | Abcam | 5 µg/ml |
| PCK | ab6410 | PCK-26 | Abcam | 5 µg/ml |
| CD20 | sc-393894 | D-10 | SCBT | 5 µg/ml |
| FoxP3 | 74816SF | D2W8E | CST | 5 µg/ml |
| NaKATPase | ab197713 | EP1845Y | Abcam | 5 µg/ml |
| CD27 | ab283697 | EPR8569 | Abcam | 5 µg/ml |
| aSMA | ab184675 | 1A4 | Abcam | 5 µg/ml |
| GATA3 | ab210672 | EPR16651 | Abcam | 5 µg/ml |
| FAP | AF3715 | polyclonal | R&D Systems | 5 µg/ml |
| HLADR | sc-53319 | TAL 1B5 | SCBT | 5 µg/ml |
| Granzyme B | ab225472 | EPR20129-217 | Abcam | 5 µg/ml |
| CD3 | ab245731 | SP162 | Abcam | 5 µg/ml |
| Vimentin | ab194719 | EPR3776 | Abcam | 5 µg/ml |
| CD56 | MA5-11563 | 56C04 | Invitrogen | 5 µg/ml |
| IgA | ab97219 | polyclonal | Abcam | 5 µg/ml |
| Thy1 | ab181885 | EPR3132 | Abcam | 5 µg/ml |
| CD11b | ab307523 | EPR1344 | Abcam | 5 µg/ml |
| PDL1 | ab215251 | EPR19759 | Abcam | 5 µg/ml |
| PDPN | 916610 | D2-40 | BioLegend | 5 µg/ml |
| CK19 | ab205445 | EPR1579Y | Abcam | 5 µg/ml |
| CD38 | ab108403 | EPR4106 | Abcam | 5 µg/ml |

